# Frontal connectivity dynamics encode contextual information during action preparation

**DOI:** 10.1101/2025.03.06.641802

**Authors:** Eleonora Arrigoni, Giacomo Guidali, Nadia Bolognini, Alberto Pisoni

**Author notes:** These authors contributed equally to this work (Eleonora Arrigoni, Giacomo Guidali). ***Corresponding authors:*** Eleonora Arrigoni, PhD – Department of Psychology, University of Milano-Bicocca, Piazza dell’Ateneo Nuovo 1, Milan, Italy;, Alberto Pisoni, PhD – Department of Psychology, University of Milano-Bicocca, Piazza dell’Ateneo Nuovo 1, Milan, Italy.

## Abstract

The context in which we perform motor acts shapes our behavior, with movement speed and accuracy modulated by contingent factors, such as the occurrence of cues that trigger or inhibit our actions. This flexibility relies on network interactions encompassing premotor and prefrontal regions, including the supplementary motor area (SMA) and the right inferior frontal gyrus (rIFG). However, the dynamic interplay between these regions during action preparation and execution based on contextual demands remains unclear. Here, we demonstrate that contextual information is encoded in SMA and rIFG interareal connectivity before action. Using Transcranial Magnetic Stimulation (TMS) and electroencephalography (EEG) during Go/No-Go tasks with varying target probabilities, we found that, during the preparatory stages of action, α-band rIFG connectivity increased in contexts where motor responses were more frequently withheld. In contrast, SMA exhibited a reversed pattern only near the target onset. Finally, β-band connectivity encoded proactive inhibition processes, increasing when action likelihood was low. Accordingly, during response implementation, both areas exhibited greater β-band connectivity when action was withheld compared to when a motor response was required, further supporting its role in inhibitory control. Our results demonstrate that α- and β-band oscillatory network dynamics support context-sensitive adaptations, illustrating how premotor and prefrontal regions synergistically modulate their interactions as they transition from preparation to response. These findings advance understanding of how the brain integrates predictive information to dynamically organize motor and cognitive resources before an action unfolds, revealing that connectivity encodes critical information driving behavior.

**Significance statement:** Adaptive behavior relies on the brain’s ability to anticipate and adjust actions based on contextual cues. Using Transcranial Magnetic Stimulation and electroencephalography (TMS-EEG), we show that the supplementary motor area (SMA) and the right inferior frontal gyrus (IFG) dynamically modulate their functional connectivity based on target predictability during action preparation and initiation. We reveal distinct oscillatory mechanisms by which SMA enhances motor readiness and IFG supports inhibitory control before action execution. These findings provide new insights into how frontal networks integrate probabilistic information to optimize action outputs in predictable contexts, with implications for disorders involving deficits in motor planning and control.

## 1. INTRODUCTION

Action anticipation, i.e., the brain’s ability to predict upcoming events and prepare for subsequent motor plans, is essential when timely and coordinated actions are crucial to adjust our behavior to meet task demands flexibly (Bogacz et al., 2010). Several prefrontal and premotor regions become active during action anticipation, linking cognition to motor output (Di Russo et al., 2017). These regions exert top-down control over the motor cortex, modulating its activity based on the anticipated action (Bestmann and Krakauer, 2015; Di Russo et al., 2017) and constituting a pre-movement “break/accelerator” system that prepares for quick and accurate responses to action triggers (Di Russo et al., 2016). Indeed, examining the temporal dynamics of pre-movement brain activity, event-related potentials (ERPs) studies identified anticipatory components originating in the supplementary motor area (SMA - Shibasaki and Hallett, 2006) and the right inferior frontal gyrus (IFG - Di Russo et al., 2016) during the early stages of action preparation (Di Russo et al., 2019). These preparatory ERPs have been linked to behavioral performance (Di Russo et al., 2016; Bianco et al., 2017), and they were shown to be influenced by probabilistic contextual information (Lucci et al., 2016). For instance, in Go/No-Go paradigms where Go trials are more likely to occur, the pre-stimulus ERP activity originating in the SMA (i.e., Bereitschaftspotential, Shibasaki and Hallett, 2006) increases, indexing enhanced motor preparation related to faster response. Conversely, when the target is rare and more cognitive control is required, there is an increase of a component originating in the right IFG (i.e., prefrontal negativity, Berchicci et al., 2012), which enhances response accuracy at the cost of longer latencies (Lucci et al., 2016).

Importantly, the SMA influences motor output through bilateral projections to the primary motor cortex (M1, Luppino et al., 1993), affecting motor response times (Welniarz et al., 2019) through a distributed network including subcortical and prefrontal regions, switching between promoting and restraining motor actions as needed (Chen et al., 2010; Picazio et al., 2014; Potgieser et al., 2014). The right IFG, instead, exerts an inhibitory effect on the contralateral M1 when movements need to be reprogrammed in response to environmental inputs (Buch et al., 2010; Neubert et al., 2010). Additionally, the right IFG’s influence on M1 varies with the predictability of the ‘go’ signal in a Go/No-Go task, being either inhibitory or facilitatory depending on the context (Campen et al., 2013).

In a previous study (Bianco et al., 2023), we explored SMA connectivity in an equiprobable Go/No-Go task. Here, we probed and recorded its activity in early and late action preparatory stages with concurrent Transcranial Magnetic Stimulation and electroencephalography recording (TMS-EEG). TMS-EEG allows to assess functional and effective connectivity of the target region by studying the amount and oscillatory profile of information that propagates from it after the TMS pulse (Thut and Miniussi, 2009; Hernandez-Pavon et al., 2023). We observed that SMA-right IFG connectivity changes throughout the pre-stimulus period, suggesting that these areas do not function in isolation but are part of a cortical network that dynamically reorganizes based on task demands (Bianco et al., 2023). However, SMA-IFG interareal dynamics during action preparation and inhibition are still poorly understood, especially concerning their adaptation to environmental demands, such as when proactive inhibitory control is required or when an action needs to take place.

Given these premises, in the present work, we hypothesized that changes in inter-regional brain communication supported by long-range phase synchronization of neuronal oscillations (Von Stein and Sarnthein, 2000; Fries, 2015) could be a critical mechanism for flexible behavioral adaptation in light of contextual information during action anticipation. To explore this, participants performed a visuomotor Go/No-Go task during TMS-EEG, where the frequency of ‘go’ target occurrences varied between experimental conditions. Here, functional connectivity of SMA and right IFG in the alpha- and beta-bands was assessed at three different time points, before and after the onset of the imperative visual stimulus.

## 2. MATERIALS and METHODS

### 2.1. Participants

Twenty-eight healthy volunteers (14 females, mean (M) age = 25.08 years, standard deviation (SD) = ±3.74) took part in the study. All participants were right-handed, as evaluated with the Edinburgh Handedness Inventory (Oldfield, 1971), had normal or corrected-to-normal vision, and no history of neurological, psychiatric, or other relevant medical conditions. Following TMS safety standards (Rossi et al., 2021), each participant filled out a safety screening questionnaire to rule out the existence of contraindications to TMS administration. After explaining the experimental procedures, all participants gave written informed consent. The study was performed in the TMS-EEG laboratory of the Department of Psychology of the University of Milano-Bicocca in accordance with the Declaration of Helsinki and following the approval of the local Ethics Committee (protocol number:701-2022). All participants were naïve to the aims of the study. Tasks’ scripts, dataset, and analysis of this study will be publicly available at Open Science Framework (OSF – https://osf.io/yteg8/)

### 2.2. Experimental Procedures

Participants performed a Go/No-Go task in which the frequency of each trial type was manipulated to create two conditions: (a) Go-Frequent (75% Go, 25% No-Go) and (b) Go-Rare (25% Go, 75% No-Go). Participants were randomly assigned to one of these conditions to avoid potential perceptual learning effects or carry-over effects of one task on the other. Hence, our sample size was divided into two groups of 14 participants each (i.e., Go-Frequent group and Go-Rare group). The number of female and male participants was balanced for each group to prevent gender-related confounding effects, as previous evidence has shown that men and women can have different action preparation styles for the same task (Bianco et al., 2020). The experimental procedure was the same for the two groups, except for the frequency of Go/No-Go trials in the experimental task.

Participants sat comfortably in a semi-reclined armchair positioned 114 cm in front of a 20" computer screen, with their arms on the armrests, ready to press a button on a PC mouse with their right-hand index finger. The experiment consisted of two recording sessions, separated by at least 48 hours, lasting about 2 hours and 30 min each. The two sessions differed only for the site of TMS administration, i.e., left SMA or right IFG. In each session, after the preparation of the EEG cap and the neuronavigation procedures, participants performed the blocks of the visual Go/No-Go task during TMS-EEG recording (Bianco et al., 2023). Before administering the Go/No-Go task, a brief practice session familiarized participants with the visual stimuli and task instructions.

### 2.3. Go/No-Go task

The Go/No-Go task used in the present study was a modified version adapted from our previous TMS-EEG study (Bianco et al., 2023). Here, TMS was delivered before (*Preparation* trials) or after (*Response* trials) the visual presentation of the Go/No-Go imperative stimulus to investigate cortical reactivity during action preparation and initiation.

Each trial of the task began with a fixation point (i.e., a yellow dot displayed on a black background), which remained at the center of the screen for the entire trial duration. In *Preparation* trials, the TMS pulse was delivered during the pre-stimulus stage after a randomly jittered interval ranging from 2900 ms to 3900 ms. Following the stimulus onset asynchrony (SOA) of either 300 ms or 700 ms, one of four different visual stimuli – gray squared configurations with black vertical and horizontal bars (4’’x4’’ of visual angle) – was randomly presented at the center of the screen for 250 ms. Hence, we delivered TMS 700 ms or 300 ms before the onset of the visual Go/No-Go target (Bianco et al., 2023). Conversely, in *Response* trials, we delivered TMS in the post-stimulus stage, 200 ms after the target presentation. At this time, the brain has generally completed the perceptual processing of the visual stimulus and is transitioning into either action initiation (for Go trials) or inhibition (for No-Go trials) (Di Russo et al., 2016). According to the experimental session, TMS was delivered over the participant’s left SMA or right IFG. Considering Go/No-Go visual stimuli, two gray squared configurations served as targets, requiring a motor response (Go trials), while the other two served as non-targets, requiring no response (No-Go trials; **Figure 1**). Participants were instructed to press the left key of the PC mouse as soon as a target stimulus (i.e., Go trials) appeared on the screen. Responses were collected within a maximum of 1500 ms from visual stimulus presentation.

**Figure 1.**
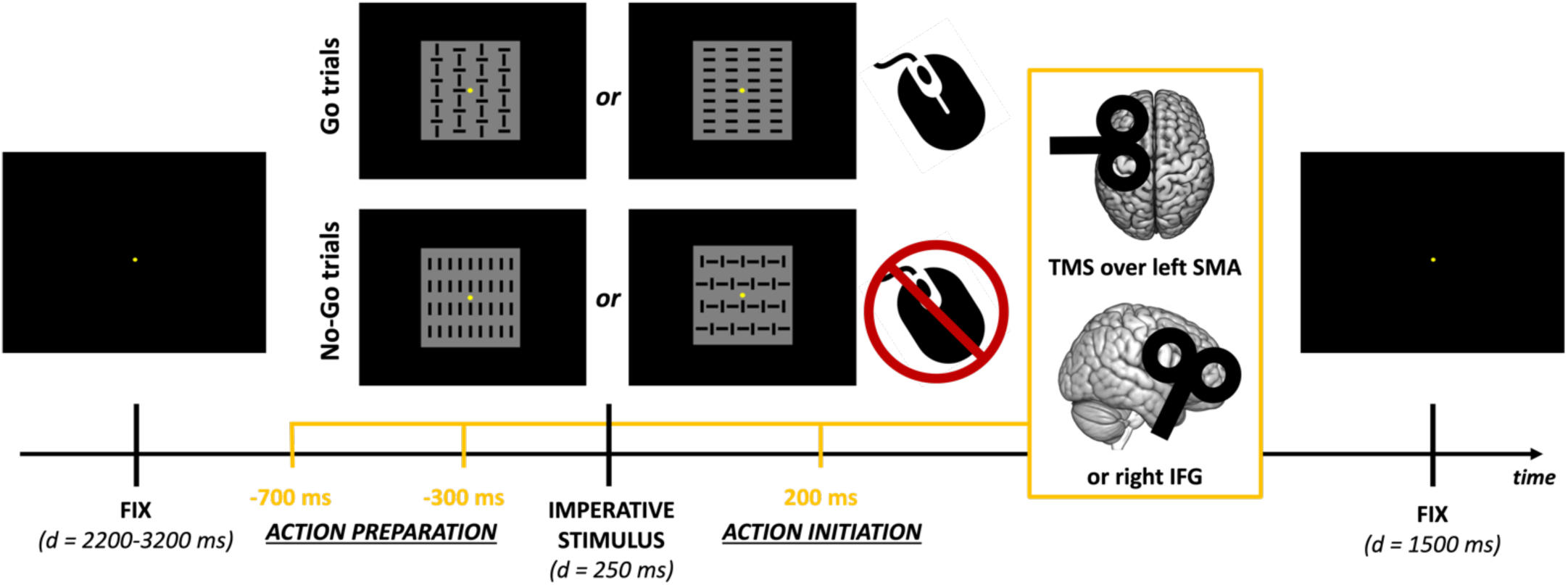
Go/No-Go task. Trials started with a fixation screen, presented for a jittering interval between 2900 ms and 3900 ms. Within this interval, TMS was delivered −700, −300 ms before or +200 ms after instruction stimulus onset on the left SMA or right IFG region. The stimulus appeared for 250 ms. Go trials appeared with 75% probability in the Go-Frequent group and 25% probability in the Go-Rare group. Each figure appeared in 50% of the Go or No-Go trials. Finally, a fixation screen was displayed to collect the participant’s response for 1500 ms.

A total of 576 trials were presented to participants throughout an experimental session. Considering the Go-Frequent group, 432 trials depicted Go visual stimuli (75%), and 144 depicted No-Go ones (25%). These numbers were the opposite for the Go-Rare group (i.e., 144 Go, 432 No-Go trials). We divided the task into 4 blocks of 144 trials to allow for a short break and avoid excessive fatigue for the participant. Within a single block, 48 trials were randomly delivered for the three SOAs (i.e., −700, −300, +200 ms). Each block lasted approximately 10 minutes, with their order counterbalanced between participants. The stimuli and timing of TMS were under computer control (E-Prime 2.0, Psychology Software Tool, Inc.).

### 2.4. TMS Parameters

Biphasic TMS pulses were delivered using an Eximia TMS stimulator (Nexstim, Helsinki, Finland) with a figure-of-eight coil (diameter = 60 mm). Stimulation sites (i.e., left SMA and right IFG) were localized on individual, high-resolution (1 mm^3^) MRI images. We selected the coordinates of the left SMA (MNI: X=-4, Y=17, Z=65) and the right IFG (X=60, Y=15, Z=27) based on previous literature on action preparation (e.g. Di Russo et al., 2016). These coordinates were used to aid experimenters in correctly identifying the stimulation hotspots on the participant’s MRI. Neuronavigation procedures were carried out using a Navigated Brain Stimulation (NBS) system (Nexstim) with infrared-based frameless stereotaxy, allowing precise monitoring of coil position and orientation and real-time estimation of the intracranial electric field’s distribution and intensity (V/m). TMS intensity and coil orientation were adjusted for each participant and stimulation site before administering the Go/No-Go task to minimize muscular artifacts and ensure the recording of early cortical response of at least 6 μV in the early components (Bianco et al., 2023). The mean induced electric field intensity was (mean ± SD): 98.21 ± 11.4 V/m for left SMA stimulation (with no differences between groups: *t*_26_ = −0.82, *p* = .42) and 100.36 ± 15.63 V/m for right IFG one (with no differences between groups: *t*_26_ = 0.84, *p* = .41), corresponding to a maximum stimulator output intensity of 55.28% (SD = 5.99) and 54.14% (SD = 6.9) for left SMA and right IFG stimulation, respectively.

### 2.5. EEG Recording and Preprocessing

EEG data were continuously acquired from a 60-channel EEG cap (EasyCap, BrainProducts GmbH, Munich, Germany) using a sample-and-hold TMS-compatible system (Nexstim™, Helsinki, Finland). Two electrodes were placed on the forehead as ground and reference. Two additional channels were placed near the eyes (one above the right eyebrow and the other on the left cheekbone) to detect ocular artifacts from eye movements and blinking. Noise masking was performed by continuously playing an audio track through earplugs, created by shuffling TMS discharge noise to prevent auditory evoked potentials (Bianco et al., 2023; Guidali et al., 2025). The volume of the noise masking was individually adjusted before each session to cover TMS clicks fully. Electrodes’ impedance was tested before each experimental session and kept below 5 kΩ. EEG signals were acquired with a sampling rate of 1450 Hz.

EEG preprocessing was carried out in MATLAB (MathWorks, Natick, MA, USA) using EEGLAB (Delorme and Makeig, 2004) and TESA toolbox (Rogasch et al., 2017) functions. Data were downsampled to 725 Hz and re-referenced using an average reference, segmented in epochs starting 1000 ms pre- and ending 1000 ms post-TMS pulse, and baseline-corrected between −500 and −250 ms before TMS pulse to avoid baseline contamination with the visual-evoked activity in the *Response* trials. Then, single trials with excessive artifacts were rejected by visual inspection. We applied the source-estimate-utilizing noise-discarding algorithm (SOUND, see Mutanen et al., 2018) implemented in TESA to attenuate extracranial noise coming from bad channels, exploiting a 3-layer spherical model with default parameters (λ = 0.1, as in Mutanen et al., 2018). Then, Independent Component Analysis (FastICA, pop_tesa_fastica, ‘tanh’ contrast) was performed after Principal Component Analysis (PCA) compression to 30 components (pop_tesa_pcacompress) to remove blinks, eye movements, residual electrical artifacts, and spontaneous muscular activity (Hernandez-Pavon et al., 2012). We further applied a semiautomatic signal space projection method for muscle artifact removal (SSP-SIR) to suppress TMS-evoked muscle artifacts post-TMS (Mutanen et al., 2016). Epochs were band-pass filtered from 1 to 80 Hz and band-stop filtered from 48 to 52 Hz using a 4th-order Butterworth filter. Finally, trials were split by condition and prepared for source reconstruction, connectivity estimation, and statistical analysis. Grand averages of TEPs obtained by stimulating the left SMA and right IFG across experimental conditions are presented in **Figures S1** and **S2**.

### 2.6. Quantification and statistical analysis

All statistical analyses were carried out using Matlab (2019b) and Jamovi (version 2.5.3; The Jamovi Project, 2025). Significant interactions were explored in mixed models with Scheffé post-hoc comparisons and repeated measures analysis of variance (rm-ANOVA) with multiple Tukey HSD post-hoc comparisons. Partial eta-squared (η_p_^2^) and Cohen’s *d* were calculated in every rmANOVA and t-test, respectively, and reported as effect size values. If not otherwise specified, mean ± standard error is reported in the **Results** section for each variable.

#### 2.6.1. Behavioral Data Analysis

We conducted a series of statistical analyses on reaction times (RTs) and accuracy to verify that the manipulation of trial probability modulated expectancy of the upcoming stimulus, affecting participants’ response behavior in the expected direction (i.e., faster RTs for the Go-Frequent group, slower RTs for the Go-Rare group).

RTs of correct Go trials were filtered within ±3 SD to account for outliers. After checking for normality violations through the Q-Q plot and skewness/kurtosis assessment, we applied base-ten logarithmic transformation on the raw RTs [i.e., log_10_(RTs)] to make the distribution closer to normality. The log-transformed RTs were then compared across experimental conditions with a Linear Mixed Model, using the ‘lmer’ model implemented in the GAMLj module in Jamovi (Gallucci, 2019). We considered ‘Group’ (Go-Frequent, Go-Rare), ‘SOA’ (−700ms, −300ms, +200 ms), and their interactions as fixed effects. We included by-subjects random intercepts to account for inter-individual differences. To account for potential confounds (e.g., fatigue, general level of arousal, learning effects) arising from the fact that TMS was delivered to the two cortical targets on separate days, we included the ‘TMS target’ factor (left SMA, right IFG) as a by-subjects random slope in our model.

Accuracy was compared across conditions using the Generalized Mixed Models for binomially distributed variables with the procedure implemented in GAMLj. Similar to RTs analysis, we considered ‘Group’, ‘SOA’, ‘TMS target’, and their interactions as fixed effects. By-subjects random intercepts and ‘TMS target’ random slope were also included in the model, and the blocks’ order was considered as a covariate.

#### 2.6.2. Electrophysiological Data Analysis

##### 2.6.2.1. Source reconstruction

Source reconstruction followed the same pipeline as described in (Bianco et al., 2023; Guidali et al., 2025). In detail, TMS-evoked activity was reconstructed at the source level to evaluate the effective connectivity of the left SMA and right IFG at the different stages of action preparation and initiation. The forward model was created starting from a Boundary Element Model (BEM) obtained by segmenting a subject MRI into five standard tissues (Gray and white matter, CSF, Skull, and Scalp). The head model was created by assigning standard conductivity values for the scalp, skull, and brain compartments (Fuchs et al., 2002; Vorwerk et al., 2014). Cortical reconstruction and volumetric segmentation of the grey matter was performed in Freesurfer (Fischl, 2012), downsampled to 8194 cortical sources, and realigned to the head model space. Individual lead-field matrices were created by aligning the forward model with the individual digitized electrode positions. eLORETA (Pascual-Marqui et al., 2011) was used for the inverse solution. EEG source reconstructed time series were collapsed into 88 regions of the AAL atlas (Tzourio-Mazoyer et al., 2002) and averaged for each cortical parcel.

##### 2.6.2.2. Connectivity analysis

We performed time-frequency decomposition with a multitaper method implemented in Fieldtrip (Oostenveld et al., 2011). Then, we computed the debiased weighted Phase Lag Index (wPLI, Vinck et al., 2011) between the two cortical parcels of the AAL atlas corresponding to the cortical sites that were targeted by TMS (i.e., the left SMA and the right IFG – pars opercularis) and the other brain parcels for alpha (8-12 Hz) and beta (13-30 Hz) bands for each TMS SOA. To account for potential spurious connectivity results, we performed the same analysis on a surrogate dataset and statistically compared it with the original data. In detail, for each participant and each condition, surrogate data were created by shuffling the phase of the source reconstructed time series of real data for each experimental condition using the ‘‘phaseran’’ Matlab function to randomize the phase while keeping the oscillatory power comparable to the original. The surrogate dataset was then entered into the same connectivity analysis pipeline. Finally, real data were compared with the surrogate ones through a permutation t-test performed on each connectivity pair. The test was corrected for multiple comparisons based on 10000 permutations with a significance level of *p* = .05 (Pisoni et al., 2018; Bianco et al., 2023; Guidali et al., 2025). Surviving connections were visualized on a surface template to highlight the resulting connectivity of the two TMS targets. Finally, the *connectivity strength* index (Guidali et al., 2025) was computed for each participant as the sum of all significant connections, allowing us to quantify the global functional connectivity evoked by TMS over our target regions. Importantly, we run separate statistical analyses for *Preparation* and *Response* trials to distinguish clearly between the neural processes involved in action preparation and anticipation and those involved in response execution or inhibition. This would also avoid possible confounds related to sensory processing (i.e., present only in post-stimulus trials).

First, we analyzed the time course of left SMA and right IFG connectivity during the pre-stimulus stage of the task, i.e., in *Preparation* trials (SOA: −700, −300 ms). This was done to determine whether the modulation of target expectancy resulted in a differential specialization of interregional communication between these two areas and the rest of the brain before stimulus presentation, depending on the task demands. To this aim, we run an rm-ANOVA on the *connectivity strength* values with the between-subjects factor ‘Group’ and the within-subjects factors ‘SOA’ and ‘Area’. This analysis was conducted separately for the alpha- and beta-bands.

Then, for our post-stimulus analysis on *Response* trials (SOA: +200 ms), we focused on the more frequent trial condition for each group – Go trials for the Go-Frequent group *vs.* No-Go trials for the Go-Rare group. This was done because the more frequent trial conditions reflected the predominant task context each group experienced, providing more precise insights into the neural processes underlying each group’s primary task demand (i.e., action execution vs withholding). Moreover, analyzing the more frequent trial condition ensured an adequate number of trials for each condition (>100), warranting the robustness of the analysis. Here, we compared *connectivity strength* values through a ‘Trial’ (Go, No-Go; between-subjects) X ‘Area’ (left SMA, right IFG; within-subjects) rm-ANOVA for each frequency band to characterize the connectivity of the two regions of interest during the initial stages of response execution or inhibition.

## 3. RESULTS

### 3.1. Behavioral results

Statistical analysis of log_10_(RTs) (**Figure 2a**) showed a significant main effect of ‘Group’ (*F*_1,25.9_ = 4.81, *p* = .037), indicating that participants in the Go-Frequent group responded faster (2.67 ± 0.01) than the ones in the Go-Rare group (2.71 ± 0.01). We found a significant main effect of ‘SOA’ (*F*_2,14951.7_ = 62.74, *p* < .001): while the RTs were comparable in trials where TMS was delivered in the *Preparation* stage (i.e., at −700 ms and - 300 ms SOAs, *t*_14952.9_ = 0.36, *p_Scheffe_* = .93), response latencies were higher at SOA +200 ms (2.7 ± 0.01) compared to −700 ms (2.69 ± 0.01; *t*_14952.9_ = 9.53, *p_Scheffe_* < .001) and – 300 ms (2.69 ± 0.01; *t*_14952.9_ = 9.88, *p_Scheffe_* < .001) SOAs. Although the ‘Group’ x ‘TMS target’ interaction (*F*_1,25.6_ = 5.74, *p* = .02) resulted statistically significant, post-hoc comparisons did not reveal significant differences among groups (all *t*s < 2.23; all *p*s > .05).

**Figure 2.**
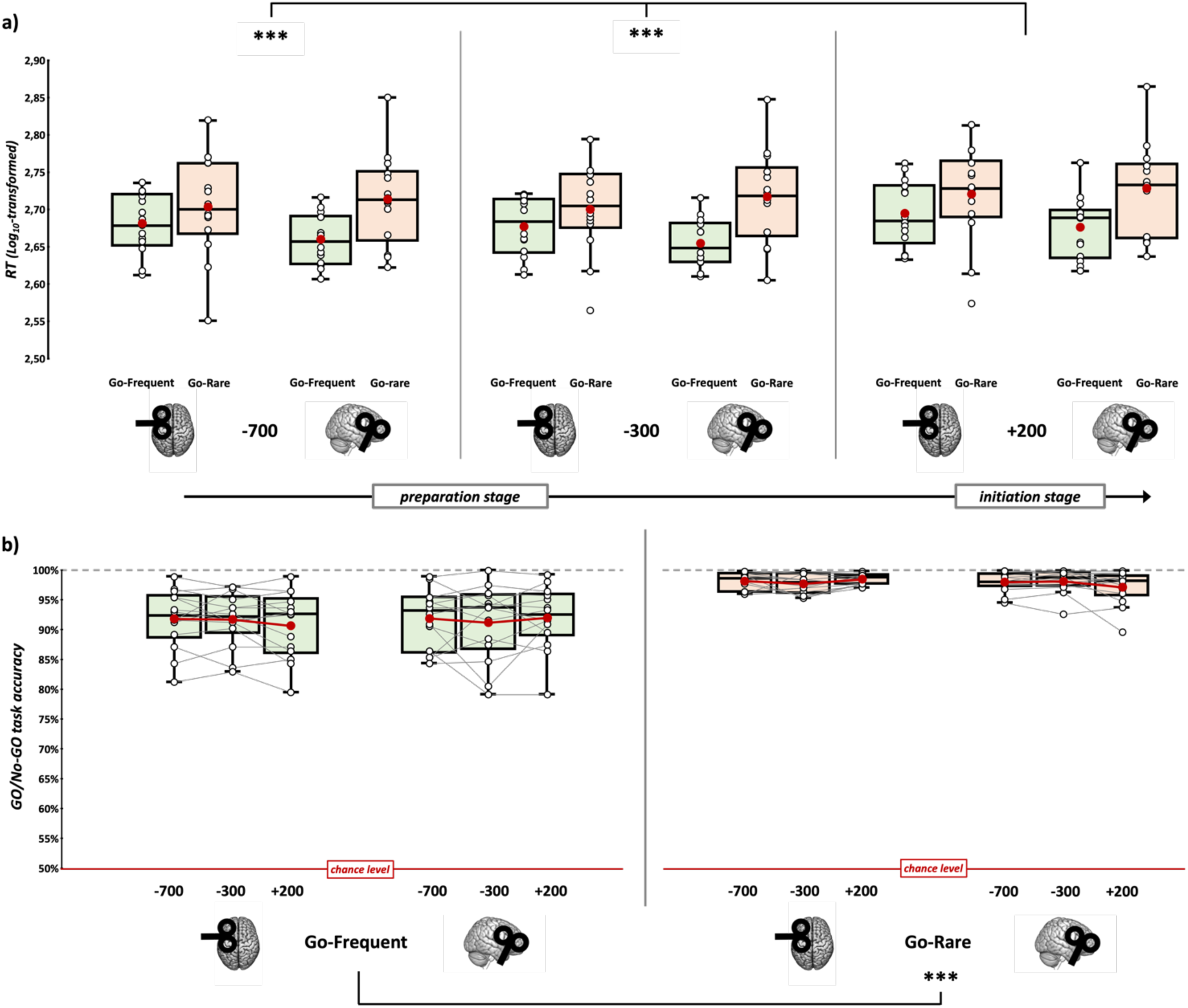
Log-transformed RTs (**a**) and accuracy (**b**) in the Go/No-Go task across experimental conditions (green boxplots = Go-frequent; orange boxplots = Go-rare). In the box-and-whiskers plots, red dots and lines indicate the means of the distributions. The center line denotes their median values. White dots and grey lines show individual participants’ scores. The box contains the 25th to 75th percentiles of the dataset. Whiskers extend to the largest observation falling within the 1.5 * inter-quartile range from the first/third quartile. Asterisks indicate significant p-values of Tukey HSD corrected post-hoc comparisons (*** = p < .001).

The main effect of ‘TMS target’ and the interactions ‘‘Group’ X ‘SOA’, ‘SOA’ X ‘TMS target’ and ‘Group’ X ‘SOA’ X ‘TMS target’ did not reach the significance level (all *F*s < 0.75, all *p*s > .05; **Table 1**).

**Table 1.** Results of the linear mixed model on the RT data across experimental conditions. The table reports each predictor’s effects and statistical significance based on Satterthwaite’s degrees of freedom approximation. Significant effects are highlighted in bold.

| Factor | *F* | df | df (res) | *p* |
| --- | --- | --- | --- | --- |
| **Group** | **4.82** | **1** | **25.9** | **.037** |
| **SOA** | **62.74** | **2** | **14951.7** | **< .001** |
| TMS Target | 0.22 | 1 | 25.6 | .65 |
| Group* SOA | 0.71 | 2 | 14951.7 | .49 |
| **Group \* TMS Target** | **5.74** | **1** | **25.6** | **.02** |
| SOA * TMS target | 0.15 | 2 | 14951.6 | .86 |
| Group * SOA * TMS Target | 0.76 | 2 | 14951.6 | .47 |

Concerning accuracy (**Figure 2b**), all the participants’ task performance was above 50% of correct trials in both experimental groups, and the mean accuracy across all conditions was consistently above 90%, indicating a generally high level of task performance. This proves that participants have performed the task attentively, regardless of the experimental condition. Our statistical analysis showed a significant main effect of ‘Group’ (χ²_1_ = 22.22, *p* < .001), indicating that the accuracy differed significantly between the two experimental groups, with the Go-Rare group having an overall more accurate performance (98 ± 1%) compared to the Go-Frequent one (91.6 ± 1.7%). The ‘SOA’ X ‘TMS Target’ (χ²_2_ = 7.49, *p* = .024) and the ‘Group’ X ‘SOA’ X ‘TMS Target’ (χ^2^_2_ = 7.99, *p* = .02) interactions reached the significance level, but post-hoc comparisons yielded no significant difference (all *ps* > 0.05). We found no other significant effect (all χ^2^s < 2.86, all *p*s > .05; **Table 2**).

**Table 2:** Results of the generalized mixed model on the accuracy data across experimental conditions. The table reports the effects and statistical significance of each predictor. Significant effects are highlighted in bold.

| Factor | $\chi^2$ | df | $p$ |
| --- | --- | --- | --- |
| **Group** | **22.22** | **1** | **< .001** |
| SOA | 2.48 | 2 | .29 |
| TMS Target | 0.14 | 1 | .71 |
| Group * SOA | 2.64 | 2 | .27 |
| Group * TMS Target | 2.85 | 1 | .09 |
| **SOA \* TMS target** | **7.49** | **2** | **.02** |
| **Group \* SOA \* TMS Target** | **7.99** | **2** | **.02** |

### 3.2. Connectivity profiles during Preparation trials

#### 3.2.1. Alpha-band connectivity

##### 3.2.1.1. Left SMA alpha-band connectivity profile during action preparation

Overall, for *Preparation* trials, alpha-band connectivity of the left SMA showed distinct patterns depending on the target probability (**Figure 3a, left panel,** see **Table S1** reporting labels of the significant parcels found). In the Go-Frequent group, connectivity shifted from frontal to posterior regions in the late preparatory phase (i.e., SOA −300 ms) compared to the early one (i.e., SOA −700 ms). In contrast, in the Go-Rare group, there was a strengthening of connectivity with bilateral frontoparietal areas just before stimulus onset.

**Figure 3.**
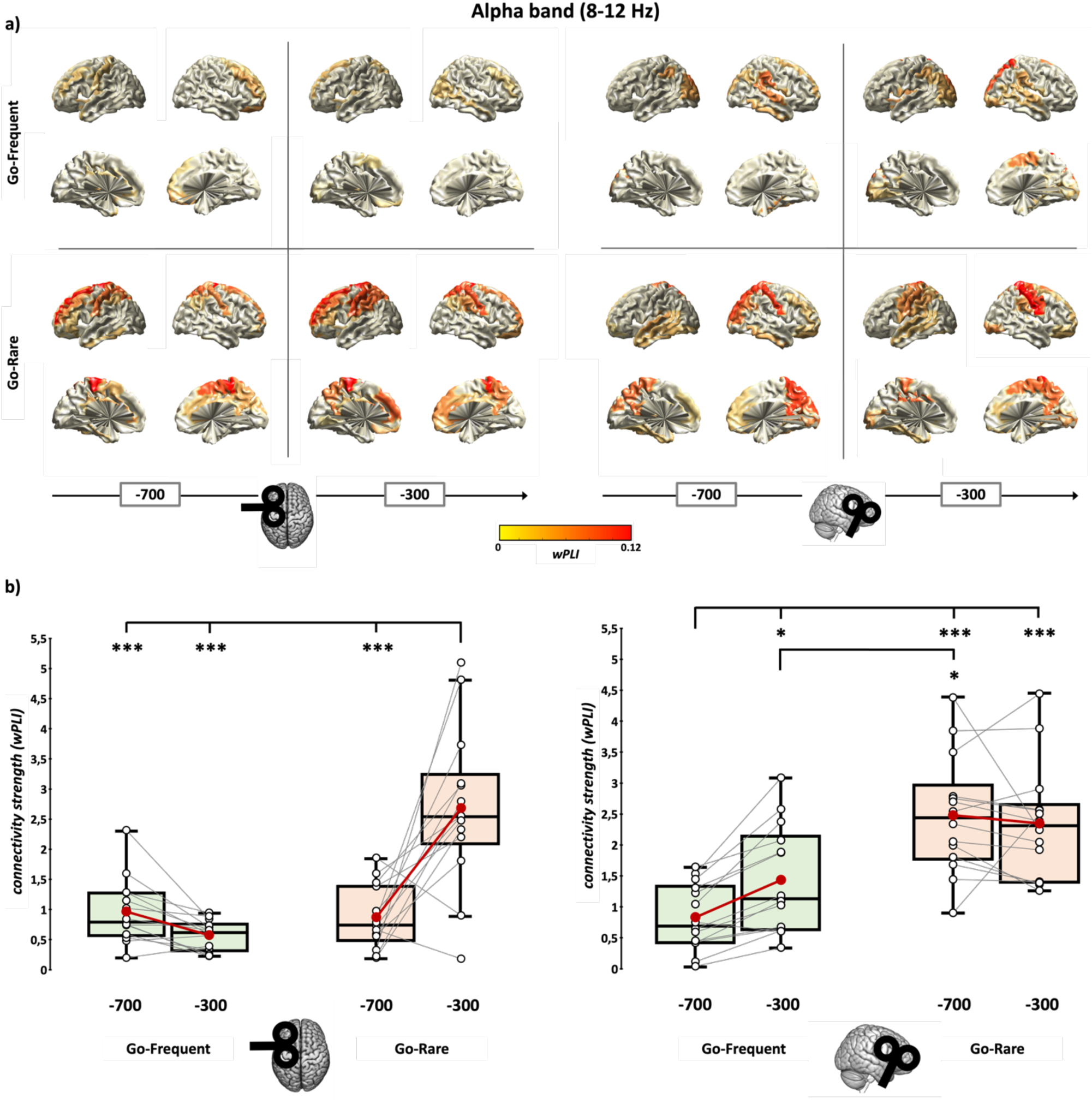
Pre-stimulus connectivity analysis across alpha (8-12 Hz) frequency band. The figure compares left SMA and right IFG functional connectivity (**a**) and the connectivity strength index values (**b**), illustrating the differences in brain network dynamics prior to stimulus onset. wPLI values of significant connections are displayed on the surface brain template for each experimental condition. TMS was triggered at an early stage (−700 ms SOA) and a later stage (−300 ms SOA) of the pre-stimulus period in each experimental group (i.e. Go-Frequent = green boxplots; Go-Rare = orange boxplots). In the box-and-whiskers plots, red dots and lines indicate the means of the distributions. The center line denotes their median values. White dots and grey lines show individual participants’ scores. The box contains the 25th to 75th percentiles of the dataset. Whiskers extend to the largest observation falling within the 1.5 * inter-quartile range from the first/third quartile. Asterisks indicate significant p-values of Tukey HSD corrected post-hoc comparisons (* = p < .05; *** = p < .001).

In the Go-Frequent group, TMS to the left SMA during preparation revealed bilateral connections with the frontal cortex, basal ganglia, and thalamus at both SOAs. SMA pre-stimulus alpha connectivity was stable with the right middle frontal gyrus, left insula, right caudate, and left thalamus, potentially indicating a preconfigured network prepared in advance to manage a habitual action with low cognitive demands and a reduced need for proactive inhibitory control. In the late preparatory period, SMA connectivity shifted to posterior regions, including bilateral occipital areas and the left superior parietal lobule.

In the Go-Rare group, the left SMA exhibited strong connections to bilateral frontoparietal areas during preparation, with significant connections to the superior and middle frontal cortex, cingulate cortex, and both superior and inferior parietal lobules detectable over time. Notably, the left SMA maintained a significant connection with the right IFG throughout the pre-stimulus phase. At SOA −300 ms, alpha-band connectivity increased towards frontal and parietal regions, including the bilateral superior and orbital frontal cortex, right insula, bilateral supramarginal gyri, left precuneus, lingual gyrus, and caudate.

##### 3.2.1.2. Right IFG alpha-band connectivity profile during action preparation

Like left SMA, right IFG alpha-band connectivity also changed over the action preparation stages (**Figure 3a, right panel, Table S2**).

Patterns from the Go-Frequent group showed that the right IFG was connected to bilateral visual and parietal areas and the right superior temporal cortex. At SOA −300 ms, connectivity increased with occipital regions. It included significant connections with the right SMA, bilateral posterior cingulate cortices, the right superior parietal cortex, medial temporal areas, and the left insula.

In contrast, in the Go-Rare group, the right IFG exhibited a stronger alpha-band connectivity pattern, linking it to the superior and middle orbital frontal regions, the cingulate cortex, the postcentral gyrus, and the precuneus in the right hemisphere, as well as bilateral visual areas, paracentral lobule, the putamen, and the left temporal cortex. In the late pre-stimulus stage (SOA −300 ms), right IFG alpha-band connectivity focused more on bilateral sensorimotor regions, connecting with bilateral M1, the right SMA, and bilateral parietal regions.

##### 3.2.1.3. Alpha-band connectivity strength changes during action preparation

The analysis of the alpha-band *connectivity strength* showed a significant ‘Group’ x ‘SOA x ‘Area’ interaction (*F*_1,26_ = 50.07, *p* < .001, η_p_^2^ = 0.66). This effect was explored by performing two separate rm-ANOVA for each area (i.e., left SMA and right IFG).

For the left SMA, we found a significant effect of ‘SOA’ (*F*_1,26_ = 11.1, *p* = .003, η_p_^2^ = 0.3), ‘Group’ (*F*_1,26_ = 26.5, *p* < .001, η_p_^2^ = 0.51) and, crucially, of the interaction ‘SOA’ X ‘Group’ (*F*_1,26_ = 26.5, *p* < .001, η_p_^2^ = 0.51). Post-hoc comparisons revealed that left SMA *connectivity strength* in – 700 ms SOA trials did not statistically differ between the two groups (Go-Frequent: 0.95 ± 0.14 *vs.* Go-Rare: 0.85 ± 0.14; *t*_26_ = 0.5, *p_Tukey_* = .96, *d* = 0.19). We did not find significant differences between −700 ms and −300 ms SOA (0.56 ± 0.07) in the Go-Frequent group (*t*_26_ = 1.29, *p_Tukey_* = .58, *d* = 0.54). In contrast, we reported an increase in *connectivity strength* between early and late (2.67 ± 0.35) preparatory phases in the Go-Rare group (*t*_26_ = 5.99, *p_Tukey_* < .001, *d* = 1.18). Finally, at −300 SOA, the left SMA exhibited stronger connectivity in the alpha-band only in the Go-Rare group (*t*_26_ = 5.89, *p_Tukey_* < .001, *d* = 2.23; **Figure 3b, left panel**).

For the right IFG, we found a main effect of ‘Group’ (*F*_1,26_ = 19.6, *p* < .001, η_p_^2^ = 0.43) and a significant ‘Group’ X ‘SOA’ (*F*_1,26_ = 8.35, *p* = .008, η_p_^2^ = 0.24) interaction. Post-hoc comparisons showed overall greater alpha-band connectivity in the Go-Rare group at −700 ms (2.46 ± 0.26 *vs.* Go-Frequent: 0.8 ± 0.14; *t*_26_ = 5.67, *p_Tukey_* < .001, *d* = 2.14) and −300 ms (2.33 ± 0.25 *vs.* Go-Frequent: 1.4 ± 0.23; *t*_26_ = 2.7, *p_Tukey_* = .049, *d* = 1.02) SOAs. *Connectivity strength* remained stable throughout the preparation period in the Go-Rare group, with no statistical differences between – 700 ms and −300 ms SOAs (*t*_26_ = 0.73, *p_Tukey_* = .88, *d* = 0.16). Conversely, we reported increased right IFG connectivity over time in the Go-Frequent group (*t*_26_ = 3.35, *p_Tukey_* = .01, *d* = 1.32; **Figure 3b, right panel**).

#### 3.2.2. Beta-band connectivity

##### 3.2.2.1. Left SMA beta-band connectivity profile during action preparation

Considering the beta-band, we found that the left SMA is primarily connected to frontoparietal regions during *Preparation* trials in both tasks (**Figure 4a, left panel, S3**).

**Figure 4.**
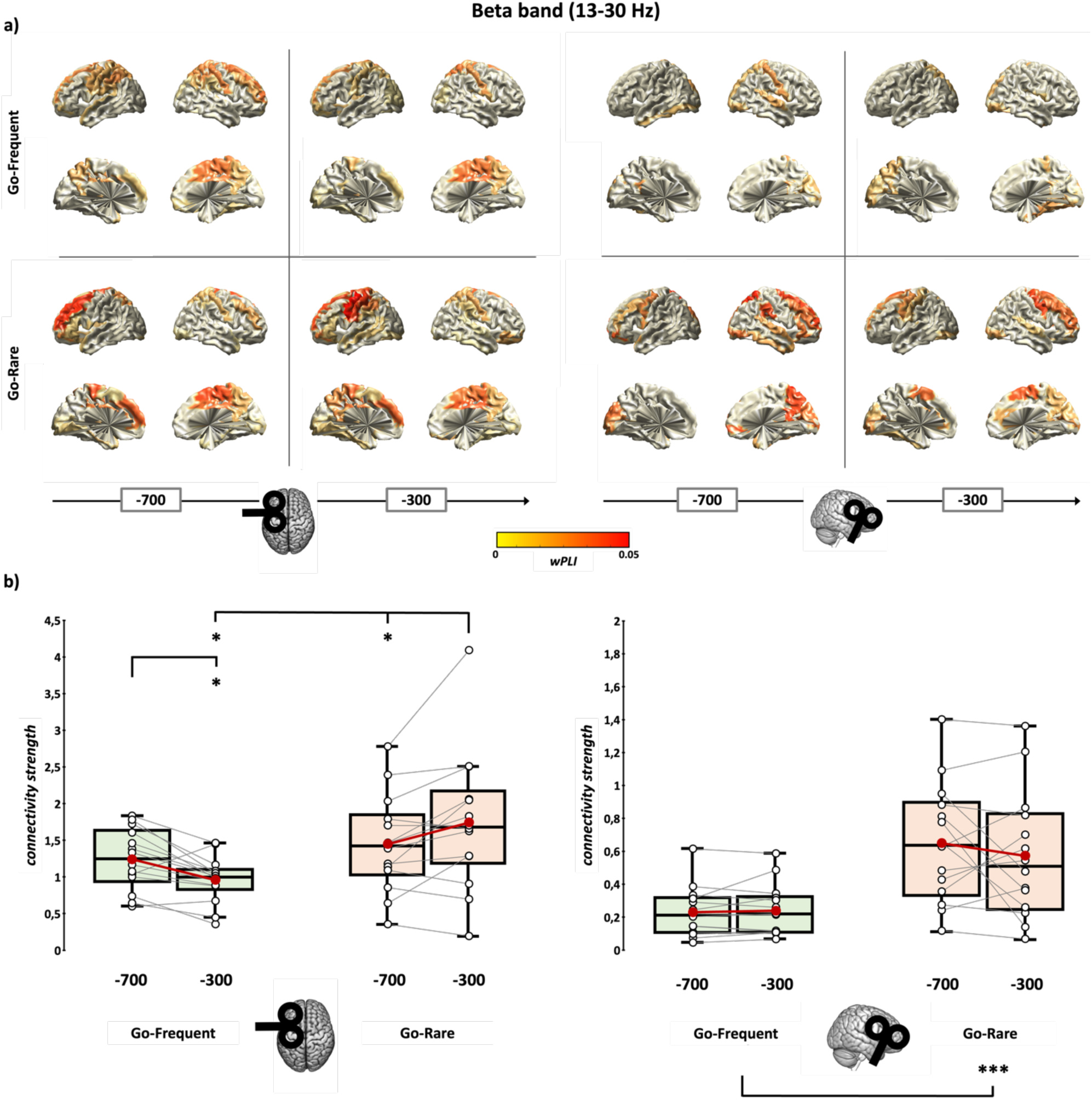
Pre-stimulus connectivity analysis across beta (13-30 Hz) frequency band. The figure compares left SMA and right IFG functional connectivity (**a**) and the connectivity strength index values (**b**), illustrating the differences in brain network dynamics before stimulus onset. wPLI values of significant connections are displayed on the surface brain template for each experimental condition. TMS was triggered at an early stage (−700 ms SOA) and a later stage (−300 ms SOA) of the pre-stimulus period in each experimental group (i.e. Go-Frequent = green boxplots; Go-Rare = orange boxplots). In the box-and-whiskers plots, red dots and lines indicate the means of the distributions. The center line denotes their median values. White dots and grey lines show individual participants’ scores. The box contains the 25th to 75th percentiles of the dataset. Whiskers extend to the largest observation falling within the 1.5 * inter-quartile range from the first/third quartile. Asterisks indicate significant p-values of Tukey HSD corrected post-hoc comparisons (* = p < .05; *** = p < .001).

During the pre-stimulus stage in the Go-Frequent group, stable connections were observed between the left SMA and the right SMA, the cingulate cortex, the left prefrontal cortex, bilateral postcentral gyri, the superior parietal cortex, the right paracentral lobule, the precuneus, and, bilaterally, the thalamus. At −700 ms SOA, the left SMA showed significant connectivity with the left M1, the right middle frontal gyrus, and bilateral parietal lobules. At −300 ms SOA, there was reduced connectivity with frontal and parietal regions, particularly in the right hemisphere, and increased connectivity with visual areas. Interestingly, the connection between the left SMA and the ipsilateral M1 was not significant in this later preparation stage.

In the Go-Rare group, beta connectivity primarily involved frontal and parietal areas, similar to the Go-Frequent group, but displayed an opposite pattern over time. Indeed, as the visual stimulus approached, the left SMA increased connectivity with the caudate and the left M1 and with bilateral occipital and parietal regions. It is noteworthy that when the target probability was low, the left SMA maintained significant functional connectivity with the right IFG throughout the preparatory period. Remarkably, we did not observe any significant connections between these two regions in the Go-Frequent group.

##### 3.2.2.2. Right IFG beta-band connectivity profile during action preparation

Probing the right IFG connectivity profile in the beta-band during *Preparation* trials revealed substantial differences between the two experimental groups (**Figure 4a, right panel, Table S4**).

In the Go-Frequent group, the right IFG was primarily connected with the right Rolandic operculum, the posterior cingulate cortex, and the right superior parietal cortex, as well as the bilateral occipital and temporal visual areas with a slight shift toward the left hemisphere between the two SOAs.

Conversely, the Go-Rare group exhibited more connections overall over the course of the pre-stimulus phase than the Go-Frequent group, particularly with bilateral frontal regions. Specifically, the right IFG showed stable connections with the left M1, the right middle frontal gyrus, the right precuneus, and bilateral occipital cortices. Over time, connectivity increased with bilateral frontal regions, including the SMA, the right precentral gyrus, the right insula, and the bilateral anterior cingulate cortex, while significant connections with temporal regions diminished.

#### 3.2.2.3. Beta-band connectivity strength changes during action preparation

We compared the connectivity strength across experimental conditions through rm-ANOVA to quantify the observed beta-band changes in the connectivity profiles of left SMA and right IFG during *Preparation* trials. We found a significant ‘SOA’ X ‘Area’ X‘Group’ interaction (*F*_1,26_ = 12.67, *p* = .001, η_p_^2^ = 0.33), further explored, as done for alpha-band analyses, with separate rm-ANOVAs for each area.

Concerning the left SMA, we found a significant effect of effect of ‘Group’ (*F*_1,26_ = 4.74, *p* = .039, η_p_^2^ = 0.15), and of the interaction ‘SOA’ X ‘Group’ (*F*_1,26_ = 16.2, *p* < .001, η_p_^2^ = 0.38). We found a significant decrease in *connectivity strength* over time between −700 ms (1.24 ± 0.11) and −300 ms SOA (0.96 ± 0.08) in the Go-Frequent group (*t*_26_ = 2.78, *p_Tukey_* = .046, *d* = 0.98). In contrast, we reported an increase in *connectivity strength* between early (1.45 ± 0.18) and late (1.74 ± 0.25) preparatory phases in the Go-Rare group (*t*_26_ = 2.91, *p_Tukey_* = .035, *d* = 0.65). At −700 SOA, *connectivity strength* did not statistically differ between groups (*t*_26_ = −0.1, *p_Tukey_* = .75, *d* = −0.38). Finally, at −300 SOA, we found stronger connectivity in the beta-band during the Go-Rare task (*t*_26_ = 2.95, *p_Tukey_* = .032, *d* = 1.12) compared to the Go-Frequent one (**Figure 4b, left panel**).

For the right IFG, we found only a main effect of ‘Group’ (*F*_1,26_ = 13.9, *p* < .001, η_p_^2^ = 0.35), indicating overall higher beta-band connectivity in the Go-Rare group (0.23 ± 0.07, vs Go-Frequent: 0.6 ± 0.07), with no statistical differences between SOAs within each experimental group (‘SOA’ X ‘Group’ interaction: *F*_1,26_ = 1.12, *p* = .3, η_p_^2^ = 0.04; **Figure 4b, right panel**).

Overall, by observing the temporal evolution of connectivity patterns of left SMA and right IFG in the beta-band, we reported significant differences in the neural activity related to *Preparation* trials between the Go-Frequent and Go-Rare groups. For the left SMA, beta-band connectivity increased over the pre-stimulus stage in the Go-Rare group, predominantly linking the stimulated region to frontal and parietal areas. At the same time, it decreased in the Go-Frequent one, being mainly restricted to visual areas. Conversely, the right IFG showed greater overall connectivity in the Go-Rare group, remaining relatively stable over the action preparation stages.

### 3.3. Connectivity profiles during Response trials

#### 3.3.1. Alpha-band connectivity during response

##### 3.3.1.1. Left SMA alpha-band connectivity profile during response

Independently from the response type that participants implemented after imperative stimulus presentation in the Go/No-Go task (i.e., Go *vs.* No-Go trials, which, considering our analysis, corresponds to the most frequent condition in each experimental group, i.e., Go trials from the Go-Frequent group and No-Go trials from the Go-Rare group, respectively – see **Statistical analysis**), TMS over the left SMA revealed an alpha-band oscillating network encompassing the right SMA, the bilateral middle frontal gyri and cingulate cortices, the left postcentral gyrus, the bilateral inferior parietal lobule, the right angular gyrus, the right occipital cortex, the right temporal cortex, and the thalamus (**Figure 5a, upper panels, Table S5**).

**Figure 5.**
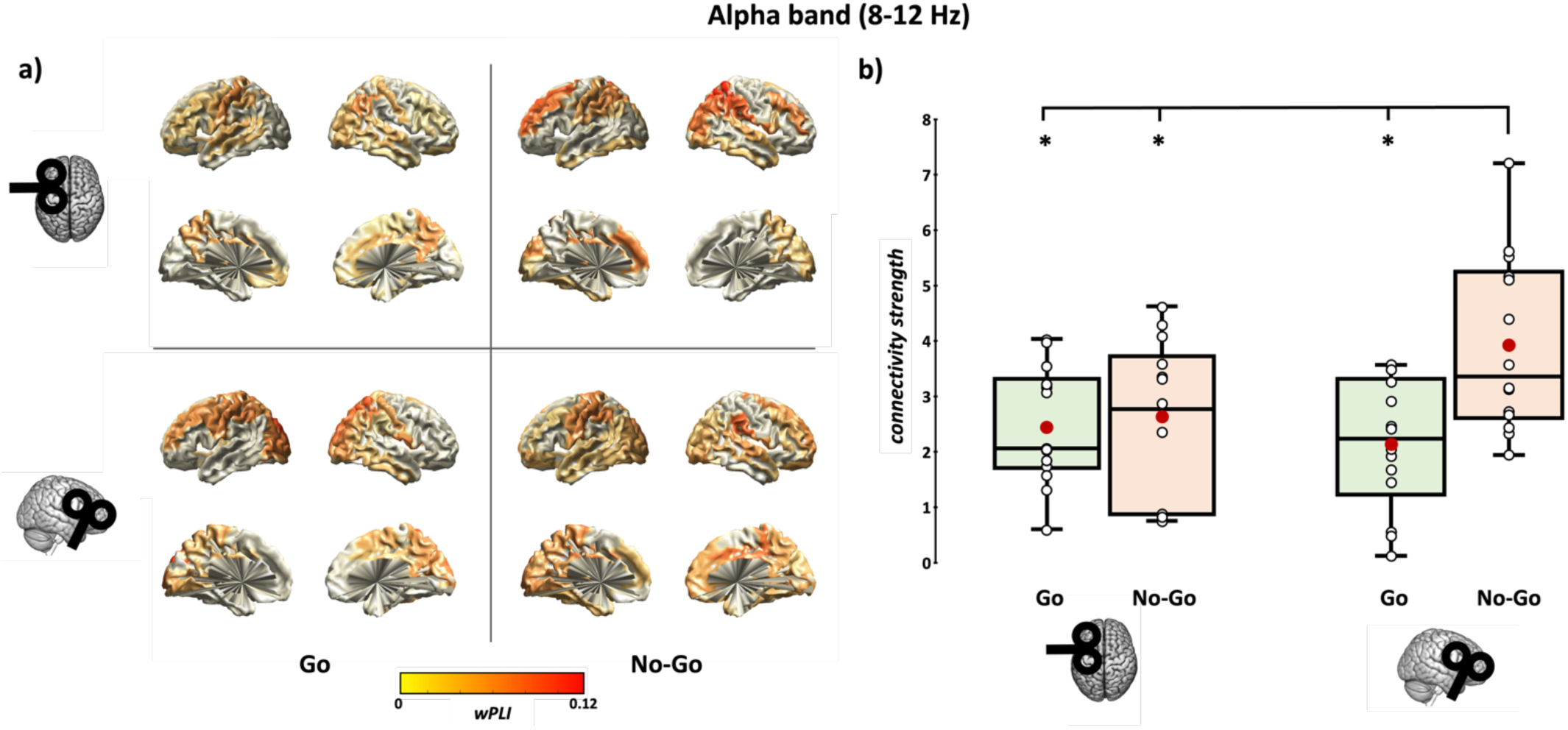
Post-stimulus connectivity analysis in the alpha (8-12 Hz) frequency band. This figure presents the changes in functional connectivity patterns for left SMA and right IFG (**a**), along with the overall connectivity strength (**b**), highlighting the TMS-evoked network response to Go trials (green boxplots) and No-Go trials (orange boxplots) in both frequency bands. wPLI values of significant connections are displayed on the surface brain template for each experimental condition. TMS was triggered at +200 ms SOA after stimulus onset in each experimental group (i.e., Go-frequent, Go trials; Go-rare, No-Go trials). In the box-and-whiskers plots, red dots indicate the means of the distributions. The center line denotes their median values. White dots show individual participants’ scores. The box contains the 25th to 75th percentiles of the dataset. Whiskers extend to the largest observation falling within the 1.5 * inter-quartile range from the first/third quartile. Asterisks indicate significant p-values of Tukey HSD corrected post-hoc comparisons (* = p < .05).

When participants implemented a motor response to the target (i.e., Go trials), the left SMA was further connected to the left M1, the left Rolandic operculum, the bilateral IFG, the bilateral orbital frontal cortex, the left middle orbital frontal gyrus, the right postcentral gyrus, and the bilateral precuneus.

Instead, when participants withheld action in response to the instructive signal (i.e., No-Go trials), alpha-band connections included the left superior frontal cortex, bilateral dorsal parietal areas, left temporal regions, and bilateral occipital regions.

##### 3.3.1.2. Right IFG alpha-band connectivity profile during response

In Go trials, the right IFG was significantly connected in the alpha-band with the left M1, left superior and middle frontal gyri, the SMA, mid-cingulum bilaterally, bilateral insula, and the right posterior cingulate cortex (**Figure 5a, lower panels, Table S5**). After the Go trials presentation, the right IFG shared strong connections with bilateral occipital and parietal regions, including the calcarine fissure, several occipital areas, postcentral gyri, and the precuneus, as well as the right putamen and right middle and inferior temporal gyri.

In No-Go trials, the right IFG exhibited a denser alpha-band connectivity profile, particularly with the bilateral frontal cortex, including the superior and middle frontal regions, and the right Rolandic operculum. It also showed significant connections with limbic structures such as the anterior cingulate cortex, medial temporal cortex, and amygdala bilaterally, along with the caudate and thalamus.

##### 3.3.1.3. Alpha-band connectivity strength during response

Comparing the alpha-band *connectivity strength* across conditions through rm-ANOVA, we found a main effect of ‘Trial’ (*F*_1,26_ = 6.72, *p* = 0.015, η_p_^2^ = 0.21), and a significant ‘Trial’ X ‘Area’ interaction (*F*_1,26_ = 6.45, *p* = .02, η_p_^2^ = 0.2). Post-hoc t-test showed that, while left SMA alpha-band *connectivity strength* did not differ between action execution (i.e., Go trials: 2.49 ± 0.28) and withholding (i.e., No-Go trials: 2.69 ± 0.37 – *t*_26_ = - 0.42, *p_Tukey_* = .97, *d* = −0.16), the right IFG connectivity was higher during response inhibition (3.96 ± 0.42, *vs.* Go trials: 2.18 ± 0.31; *t*_26_ = 3.4, *p_Tukey_* = .01, *d* = 1.29). Moreover, *connectivity strength* was significantly higher for the right IFG than for the left SMA during response inhibition (*t*_26_ = 2.88, *p_Tukey_* = .037, *d* = 0.71). At the same time, it was comparable during action execution (*t*_26_ = 0.78, *p_Tukey_* = .89, *d* = 0.21, **Figure 5b**).

#### 3.3.2. Beta-band connectivity during response

##### 3.3.2.1. Left SMA beta-band connectivity during response

During *Response* trials, in the beta-band, we found that the left SMA displayed widespread connectivity, mainly with the frontal, parietal, and occipital cortex, regardless of the decision to respond or withhold (**Figure 6a, upper panels, Table S6**). The patterns showed that while response implementation relied on a more SMA-related right-lateralized network encompassing parieto-occipital regions, action inhibition involved a more bilateral and distributed circuit.

**Figure 6.**
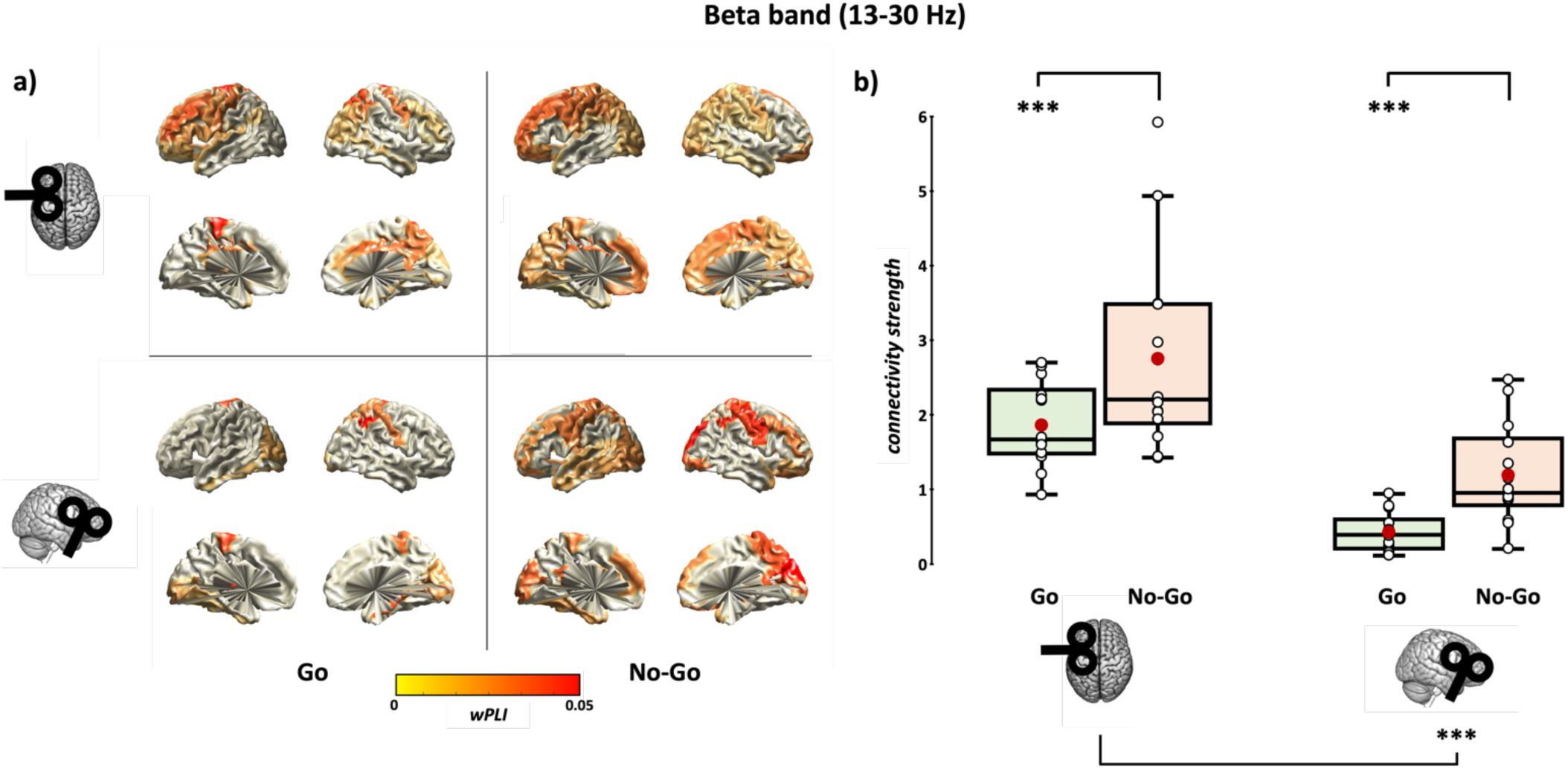
Post-stimulus connectivity analysis in the beta (13-30 Hz) frequency band. This figure presents the changes in functional connectivity for left SMA and right IFG (**a**), along with connectivity strength index values (**b**), highlighting the TMS-evoked network response to Go trials (green boxplots) and No-Go trials (orange boxplots) in both frequency bands. wPLI values of significant connections are displayed on the surface brain template for each experimental condition. TMS was triggered at +200 ms SOA after stimulus onset in each experimental group (i.e., Go-frequent, Go trials; Go-rare, No-Go trials). In the box-and-whiskers plots, red dots indicate the means of the distributions. The center line denotes their median values. White dots show individual participants’ scores. The box contains the 25th to 75th percentiles of the dataset. Whiskers extend to the largest observation falling within the 1.5 * inter-quartile range from the first/third quartile. Asterisks indicate significant p-values of Tukey HSD corrected post-hoc comparisons (*** = p < .001).

During Go trials, left SMA connectivity included bilateral precentral, IFG, superior orbital, and middle frontal regions, along with the left insula; bilateral postcentral, angular, and precuneus areas in the parietal lobe; right superior and middle occipital cortices in the occipital lobe; and the right medial temporal and amygdala, as well as bilateral caudate and thalamus, subcortically.

In No-Go trials, the left SMA displayed a more distributed network, showing increased connections with right hemisphere frontal regions, including the right superior orbital frontal cortex, the right SMA, and bilateral superior medial frontal cortex, as well as the right insula and left cingulate cortex. Additionally, the left SMA had greater connectivity with bilateral occipital and parietal regions during No-Go trials compared to the Go ones.

##### 3.3.2.2. Right IFG beta-band connectivity during response

A common pattern was found considering right IFG beta-band connectivity (**Figure 6a, lower panels, Table S6**). After stimulus presentation, irrespective of the trial type, the right IFG was connected to bilateral occipital and parietal regions.

During No-Go trials, right IFG connectivity was significantly stronger than in Go trials, with relevant connections to bilateral precentral areas, superior medial and orbital frontal cortices, middle frontal gyri, left Rolandic operculum, right orbitofrontal cortex, bilateral insula, left cingulate cortex, left medial temporal, bilateral amygdala, left superior parietal cortex, and right precuneus, as well as enhanced connectivity with the bilateral temporal cortex.

##### 3.3.2.3. Beta-band connectivity strength during response

The analysis of beta-band *connectivity strength* showed only a significant effect of the factors ‘Trial’ (*F*_1,26_ = 21.4, *p* < .001, η_p_^2^ = 0.45) and ‘Area’ (*F*_1,26_ = 37.93, *p* < .001, η_p_^2^ = 0.59). This pattern indicates that *connectivity strength* was significantly higher when participants had to withhold the motor response (No-Go trials: 1.96 ± 0.13) than when a motor response was required (Go trials: 1.13 ± 0.13). Furthermore, the left SMA presented higher *connectivity strength* in the beta-band (2.3 ± 0.19) compared to the right IFG (0.79 ± 0.1). Double interaction did not reach statistical significance (*F*_1,26_ = 0.06, *p* = .81, η_p_^2^ < 0.01; **Figure 6b**)

## 4. DISCUSSION

The main goal of the present study was to investigate cortical dynamics of the left SMA and right IFG during action preparation as a function of action anticipation. Our findings demonstrate how alpha- and beta-band connectivity of the left SMA and right IFG encode contextual information and occur in parallel to balance motor readiness and proactive inhibition synergistically.

### 4.1. Behavioral influence of target predictability on motor performance

In line with previous experiments on stimulus probability in visuomotor tasks (e.g., Dykes and Pascal, 1981; Miller and Anbar, 1981; Helton et al., 2009; Lucci et al., 2016), the Go-frequent version of the Go/No-Go task successfully facilitated motor readiness and reduced inhibition, leading to faster but less accurate motor responses. Conversely, in the Go-Rare version, targets were more difficult to predict, possibly engaging proactive inhibitory control over motor facilitation; this resulted in slower but more accurate responses. Hence, by increasing target predictability, we successfully biased participants towards motor readiness, while target uncertainty enhanced proactive inhibition to prevent premature, potentially inappropriate actions (Bestmann and Duque, 2016; Wessel and Aron, 2017). Related to these behavioral differences, we observed a modulation of cortical connectivity patterns during the *Preparation* trials.

### 4.2. Alpha-band dynamics during action preparation

During the Go-Frequent task, the left SMA was progressively more connected in the alpha band with posterior visual regions, possibly due to a bias toward the action to be implemented, leading to anticipatory recruitment of a visuomotor network (Benedetto et al., 2021), as we observed in Bianco et al. (2023). This pattern suggests that interactions between SMA and posterior regions might be crucial for top-down integration of sensory readiness and attentional, task-relevant processing in view of an anticipated stimulus requiring an already planned motor response (Capotosto et al., 2009; Palva and Palva, 2011; Sauseng et al., 2011; Lobier et al., 2018). Conversely, when target predictability was low (i.e., Go-Rare task), we found prominent alpha-band connectivity between the SMA and a widespread set of bilateral frontoparietal regions, whose strength significantly increased during the late pre-stimulus phase. Notably, left SMA is significantly connected with the contralateral IFG during the whole preparatory phase, suggesting that, when an action is difficult to anticipate, enhanced communication in the alpha band between premotor regions deputed to motor preparation and a widespread frontoparietal network may be required to compensate for stimulus uncertainty proactively. Concerning the right IFG, alpha-band connectivity in the Go-Frequent task significantly increased in the late phase, likely reflecting the progressive rise of top-down monitoring required by the task over time to avoid anticipatory responses. In the Go-Rare group, the right IFG displayed more temporally stable and widespread alpha-band connectivity, likely supporting sustained proactive inhibitory control to prevent unwanted responses. Notably, in this condition, relevant connections with the SMA and the M1 were observed in the late phase of the pre-stimulus period. These findings may indicate that interregional synchronization of alpha oscillations could be a key mechanism within prefrontal and premotor networks underlying proactive cognitive control, balancing readiness and inhibition. In the same vein, previous works suggested that alpha synchronization is crucial to aligning brain regions’ excitability, optimizing top-down interactions in large-scale networks that support sustained attention and anticipation for upcoming cognitive demands (Palva and Palva, 2011; Sadaghiani et al., 2012), particularly when the action is unpredictable.

### 4.3. Beta-band dynamics during action preparation

Considering the beta-band in the Go-Frequent condition, SMA connectivity - particularly with frontal and parietal areas, including the right IFG - decreased over time, while in the Go-Rare condition, it increased in the late preparatory stage. This finding suggests that the brain engages an early beta-band prefrontal-premotor connection to balance movement readiness when action cannot be predicted, also considering that the right IFG maintained stable beta-band connectivity in both groups, although a more distributed and stronger network was found in the Go-Rare condition.

Critically, modulations of left SMA and right IFG connections with left M1 (i.e., the region governing the effector) occurred in the preparatory period according to the probability of the target. Both left SMA and right IFG were found connected with the left M1 in the beta-band only in the Go-Rare condition, where inhibitory control is needed to maintain high control of the performance. While the left SMA-M1 connection emerges only immediately before the target presentation, communication between the right IFG and M1 is stable. These results are particularly relevant considering the role of beta oscillations in the motor system, which are typically suppressed during motor preparation and execution; this pattern marks the disinhibition of the motor system, allowing movement initiation (Neuper and Pfurtscheller, 2001; Engel and Fries, 2010; Little et al., 2019). Furthermore, beta synchronization supports effective communication between prefrontal, premotor, and motor regions, which is causally linked to motor inhibition in Go/No-go tasks (Picazio et al., 2014). Our study extended this evidence to the preparatory period, showing beta-band modulations according to task demands. Indeed, motor preparation may involve reducing premotor and prefrontal beta connectivity, while sustained beta synchronization may support proactive inhibition due to anticipation (Picazio et al., 2014; Wessel and Anderson, 2024). Another critical remark concerns the temporal dynamics of the effects: SMA connectivity demonstrated a temporal specialization throughout preparation according to action anticipation. In contrast, the right IFG did not exhibit this pattern, suggesting an all-or-nothing, ‘brake’ mechanism, possibly because the response must be suppressed until the target appearance. This may indicate that the role of the right IFG is less flexible than the influence exerted by the SMA in the preparatory period.

### 4.4. SMA and IFG communication during early stages of action implementation

We also examined connectivity patterns during *Response* trials to better interpret cortical dynamics during the preparatory phase and elucidate how alpha- and beta-band networks involving the investigated regions evolve as action is executed or inhibited.

In the alpha band, the left SMA was primarily connected to motor and sensory regions supporting motor control during execution in Go trials. Conversely, in No-Go trials, TMS over SMA highlighted a broader network, although the network strength did not significantly differ from action initiation. The right IFG, on the other hand, showed significantly more pronounced connectivity in No-Go trials, particularly with frontal regions, reflecting its stronger role in cognitive control and inhibition compared to Go trials.

In the beta-band, both the left SMA and right IFG showed broader and significantly stronger connectivity in the No-Go trials compared to the Go trials. This finding is consistent with the idea that these regions represent the main cortical hubs underlying action control, particularly for action inhibition (Wessel and Aron, 2017). However, their precise role has long been debated, questioning whether the right IFG has a functional primacy as an inhibition module or whether its inhibitory function is somehow mediated by the SMA and pre-SMA gating activity (Aron et al., 2004; Nachev et al., 2008; Hampshire et al., 2010; Swann et al., 2012; Hampshire and Sharp, 2015; Schaum et al., 2021). Our findings partly reconcile these discrepant accounts by providing a more complex network perspective.

### 4.5. Conclusions: Towards a network perspective of action preparation

The SMA and IFG cortical dynamics are engaged in complementary but distinct roles in motor response inhibition (as assessed with the Go/No-Go task) and are sensitive to contextual aspects. During the preparation phase, the SMA has a prominent role in setting a different level of motor readiness based on action anticipation, while during the response phase, the IFG is more heavily recruited in cognitive monitoring and inhibitory processes, particularly when action withholding is needed.

Our findings support a general framework in which frequency-specific oscillatory brain dynamics are crucial in flexibly preparing and implementing motor responses based on task demands. Specifically, we propose that interactions between the alpha-band activity of the SMA and the right IFG tend to be stronger when they must counter the more frequent context. Our findings provide documents on the flexible, task-dependent modulation of alpha- and beta-band connectivity of SMA and IFG during action preparation: on the one hand, alpha-band activity is potentially associated with attentional control and context detection in the right IFG (Erika-Florence et al., 2014), on the other the SMA may interact in this frequency band to modulate motor readiness to adapt to contextual demands. Conversely, SMA beta-band connectivity seems to play a critical role in balancing motor preparation and inhibition depending on the predictability of the upcoming action (Duque et al., 2013), while the right IFG interacts in this frequency to exert inhibition in a broader control network (Wessel and Anderson, 2024).

In conclusion, our study underscores the vital role of a dynamic, network-based perspective of action preparation and control. The preparation and efficient implementation of actions in response to external stimuli requires interareal interactions shaped by action anticipation well before the initiation of the motor plan, highlighting the brain’s ability to adaptively modulate its network communications in advance, depending on task demands.

## AUTHORS’ CONTRIBUTION

**Eleonora Arrigoni**: conceptualization, methodology, software, investigation, visualization, formal analysis, data curation, writing – original draft

**Giacomo Guidali**: methodology, software, investigation, visualization, formal analysis, writing – review & editing

**Nadia Bolognini**: writing - review & editing

**Alberto Pisoni**: conceptualization, methodology, supervision, resources, funding acquisition, writing – review & editing

## Acknowledgments

We thank Chiara Franciscono and Francesca Gallo for their valuable help in data collection.

