## Supplemental files for "Frontal connectivity dynamics encode contextual information during action preparation"

### - SUPPLEMENTAL MATERIALS -

This file contains the following Supplementary Materials:

- Figure S1 ………………………………… p. 2

- Figure S2 ………………………………… p. 3

-Table S1 …………………………………. p. 4

-Table S2 …………………………………. p. 7

-Table S3 …………………………………. p. 10

-Table S4 …………………………………. p. 12

-Table S5 …………………………………. p. 14

-Table S6 …………………………………. p. 17

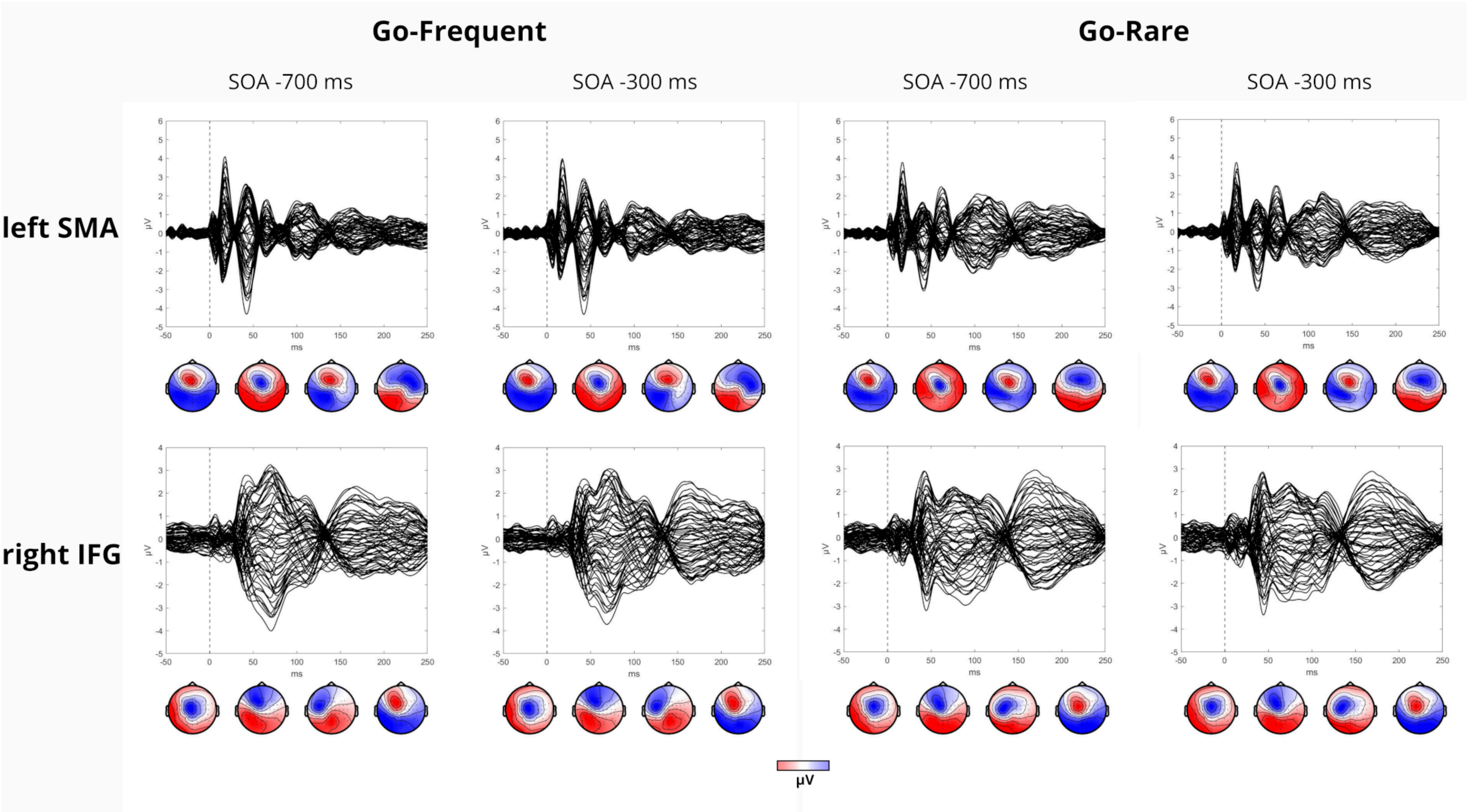


**Figure S1**. Sensor level TEP grand average and topography of the main peaks recorded during action preparation (i.e., -700 ms SOA, -300 ms SOA) in the Go-Frequent (left panels) and Go-Rare (right panels) groups during left SMA (upper panels) and right IFG (lower panels) stimulation.

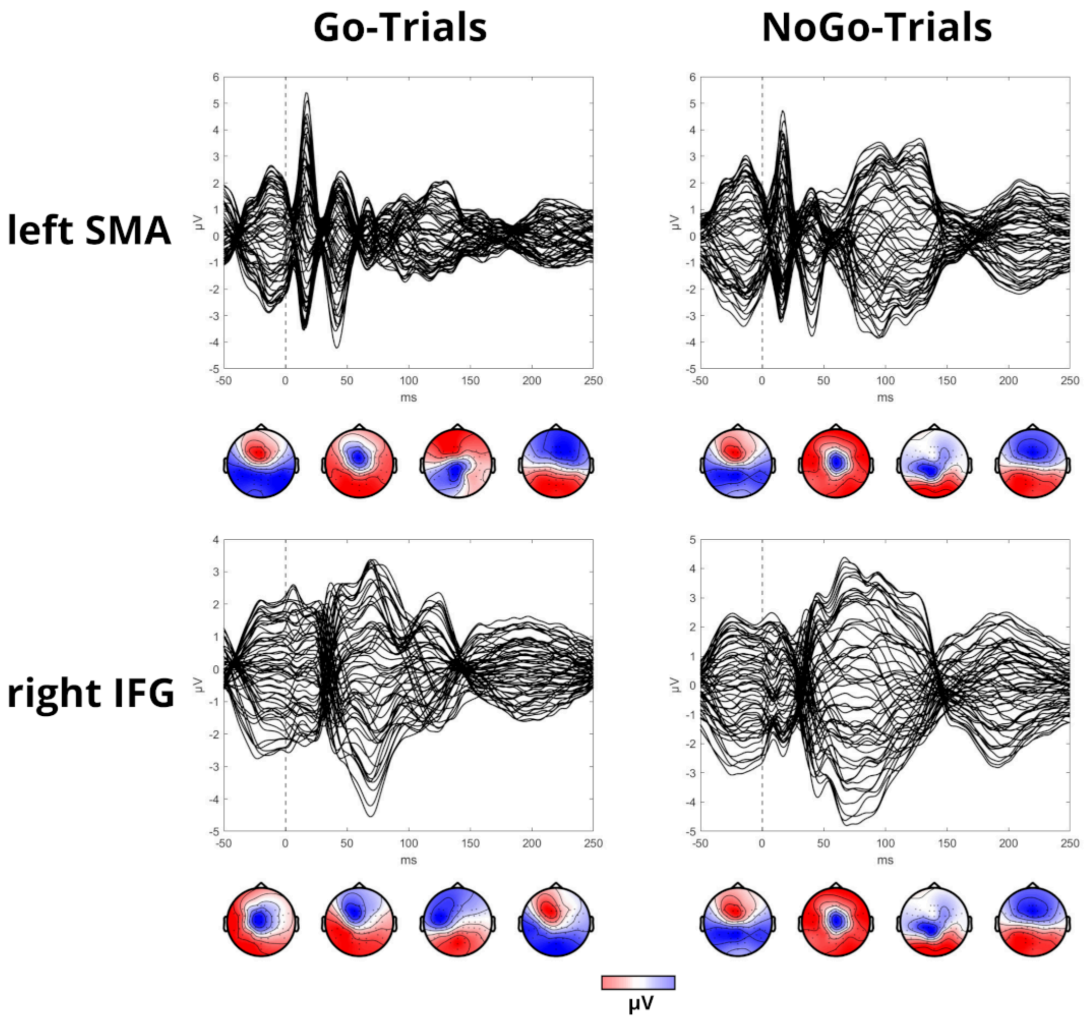


**Figure S2**. Sensor level TEP grand average and topography of the main peaks recorded during action initiation (i.e., +200 ms SOA) for GO (left panels) and No-Go (right panels) trials during left SMA (upper panels) and right IFG (lower panels) stimulation.

| ALPHA BAND (8-12 Hz) | SMA | | | |
|---|---|---|---|---|
| | GO Frequent | | GO Rare | |
| | -700 | -300 | -700 | -300 |
| Frontal Lobe | Frontal Sup Right | Frontal Sup Left | Frontal Sup Left | Frontal Sup Left |
| | Frontal Sup Orb Right | Frontal Mid Right | Frontal Sup Right | Frontal Sup Right |
| | Frontal Mid Left | Rolandic Oper Right | Frontal Sup Orb Left | Frontal Sup Orb Left |
| | Frontal Mid Right | Rectus Left | Frontal Mid Left | Frontal Sup Orb Right |
| | Olfactory Left | Insula Left | Frontal Mid Orb Right | Frontal Mid Left |
| | Olfactory Right | Cingulum Post Right | Frontal Inf Oper Right | Frontal Mid Orb Right |
| | Frontal Sup Medial Right | | Frontal Inf Tri Right | Frontal Inf Oper Right |
| | Frontal Med Orb Right | | Rolandic Oper Right | Olfactory Right |
| | Rectus Right | | Supp Motor Area Right | Frontal Sup Medial Left |
| | Insula Left | | Olfactory Right | Frontal Sup Medial Right |
| | Cingulum Ant Left | | Cingulum Ant Left | Frontal Med Orb Right |
| | Cingulum Mid Left | | Cingulum Ant Right | Rectus Left |
| | | | Cingulum Mid Left | Rectus Right |
| | | | Cingulum Mid Right | Insula Left |
| | | | Cingulum Post Left | Insula Right |
| | | | Cingulum Post Right | Cingulum Ant Left |
| | | | | Cingulum Ant Right |
| | | | | Cingulum Mid Left |
| | | | | Cingulum Mid Right |
| | | | | Cingulum Post Left |
| | | | | Cingulum Post Right |
| Temporal Lobe | Temporal Pole Mid Left | Amygdala Left | Hippocampus Left | |
| | | Temporal Mid Right | ParaHippocampal Left | |
| | | | Amygdala Left | |
| | | | Cuneus Right | |
| Parietal Lobe | Postcentral Left | Parietal Sup Left | Postcentral Left | Postcentral Left |
| | | | Postcentral Right | Postcentral Right |
| | | | Parietal Sup Left | Parietal Sup Left |

| | | | | |
|---|---|---|---|---|
| | | | Parietal Sup Right | Parietal Sup Right |
| | | | Parietal Inf Left | Parietal Inf Left |
| | | | Parietal Inf Right | Parietal Inf Right |
| | | | Angular Right | SupraMarginal Left |
| | | | Precuneus Right | SupraMarginal Right |
| | | | Paracentral Lobule Left | Angular Right |
| | | | Paracentral Lobule Right | Precuneus Left |
| | | | | Precuneus Right |
| | | | | Paracentral Lobule Left |
| | | | | Paracentral Lobule Right |
| **Occipital Lobe** | | Calcarine Left | Cuneus Right | Lingual Left |
| | | Occipital Sup Right | | |
| | | Occipital Mid Right | | |
| **Subcortical structures** | Caudate Left | Caudate Right | Caudate Right | Caudate Left |
| | Caudate Right | Thalamus Left | Putamen Right | Caudate Right |
| | Putamen Right | Thalamus Right | Temporal Inf Left | Putamen Right |
| | Thalamus Left | | | |

**Table S1.** AAL labels of the significant parcels ($p < .05$) found for the alpha band during SMA stimulation for Go Frequent and Go Rare groups for action preparation trials (i.e., -700 ms SOA, -300 ms SOA).

| ALPHA BAND (8-12 Hz) | IFG | | | |
|---|---|---|---|---|
| | GO Frequent | | GO Rare | |
| | -700 | -300 | -700 | -300 |
| Frontal Lobe | Frontal Sup Right | Rolandic Oper Left | Frontal Sup Orb Left | Precentral Left |
| | Rolandic Oper Right | Rolandic Oper Right | Frontal Sup Orb Right | Precentral Right |
| | | Supp Motor Area Right | Frontal Mid Right | Frontal Sup Right |
| | | Insula Left | Frontal Mid Orb Left | Frontal Sup Orb Right |
| | | Cingulum Post Left | Frontal Mid Orb Right | Frontal Mid Orb Right |
| | | Cingulum Post Right | Frontal Inf Orb Right | Frontal Inf Orb Right |
| | | | Olfactory Right | Rolandic Oper Left |
| | | | Frontal Sup Medial Left | Supp Motor Area Right |
| | | | Frontal Sup Medial Right | Olfactory Left |
| | | | Frontal Med Orb Right | Rectus Right |
| | | | Rectus Right | Cingulum Ant Right |
| | | | Cingulum Ant Left | Cingulum Mid Left |
| | | | Cingulum Ant Right | Cingulum Post Right |
| | | | Cingulum Post Left | |
| | | | Cingulum Post Right | |
| Temporal Lobe | ParaHippocampal Right | Hippocampus Right | Hippocampus Left | Hippocampus Left |
| | Heschl Right | ParaHippocampal Right | ParaHippocampal Left | Hippocampus Right |
| | Temporal Sup Right | Heschl Left | Amygdala Left | ParaHippocampal Left |
| | Temporal Inf Right | Heschl Right | Heschl Left | ParaHippocampal Right |
| | | Temporal Sup Right | Temporal Sup Left | Heschl Left |
| | | Temporal Mid Right | Temporal Pole Sup Left | Heschl Right |
| | | | Temporal Mid Left | Temporal Sup Left |
| | | | Temporal Pole Mid Left | Temporal Mid Left |
| | | | Temporal Inf Left | Temporal Pole Mid Left |
| | | | Temporal Inf Right | |
| Parietal Lobe | Parietal Inf Left | Parietal Sup Right | Postcentral Right | Postcentral Left |
| | SupraMarginal Right | Parietal Inf Left | Parietal Sup Right | Postcentral Right |
| | Angular Right | SupraMarginal Right | Angular Right | Parietal Sup Left |

| | | | | |
|---|---|---|---|---|
| | | Angular Left | Precuneus Left | Parietal Inf Left |
| | | | Precuneus Right | SupraMarginal Left |
| | | | Paracentral Lobule Left | SupraMarginal Right |
| | | | | Precuneus Right |
| | | | | Paracentral Lobule Left |
| | | | | Paracentral Lobule Right |
| **Occipital Lobe** | Occipital Sup Left | Calcarine Left | Calcarine Right | Calcarine Left |
| | Occipital Sup Right | Calcarine Right | Cuneus Left | Calcarine Right |
| | Occipital Mid Left | Lingual Left | Cuneus Right | Lingual Right |
| | | Lingual Right | Lingual Right | Occipital Inf Right |
| | | Occipital Sup Left | Occipital Sup Right | Fusiform Left |
| | | Occipital Sup Right | Occipital Mid Right | |
| | | Occipital Mid Left | Occipital Inf Left | |
| | | Occipital Inf Left | Fusiform Left | |
| | | Occipital Inf Right | | |
| **Subcortical structures** | Thalamus Right | | Putamen Right | Caudate Right |
| | | | Thalamus Left | Putamen Right |

**Table S2.** AAL labels of the significant parcels ($p < .05$) found for the alpha band during IFG stimulation for Go Frequent and Go Rare groups for action preparation trials (i.e., -700 ms SOA, -300 ms SOA).

| BETA BAND (13-30 Hz) | SMA | | | |
|---|---|---|---|---|
| | **GO Frequent** | | **GO Rare** | |
| | **-700** | **-300** | **-700** | **-300** |
| **Frontal Lobe** | Precentral Left | Frontal Sup Left | Frontal Sup Left | Precentral Left |
| | Frontal Sup Left | Frontal Sup Orb Left | Frontal Mid Left | Frontal Sup Left |
| | Frontal Sup Right | Frontal Mid Orb Left | Frontal Mid Right | Frontal Sup Orb Right |
| | Frontal Mid Right | Rolandic Oper Left | Frontal Mid Orb Left | Frontal Mid Orb Left |
| | Frontal Mid Orb Left | Supp Motor Area Right | Frontal Inf Oper Right | Frontal Inf Oper Right |
| | Supp Motor Area Right | Frontal Sup Medial Left | Frontal Inf Tri Right | Frontal Inf Tri Right |
| | Frontal Sup Medial Left | Insula Left | Frontal Inf Orb Right | Frontal Inf Orb Left |
| | Frontal Med Orb Right | Cingulum Mid Right | Rolandic Oper Right | Rolandic Oper Right |
| | Rectus Right | Cingulum Post Left | Supp Motor Area Right | Supp Motor Area Right |
| | Cingulum Mid Left | Cingulum Post Right | Olfactory Right | Olfactory Left |
| | Cingulum Mid Right | | Frontal Sup Medial Left | Olfactory Right |
| | Cingulum Post Right | | Insula Left | Frontal Sup Medial Left |
| | | | Insula Right | Rectus Right |
| | | | Cingulum Ant Left | Insula Right |
| | | | Cingulum Mid Left | Cingulum Mid Left |
| | | | Cingulum Mid Right | Cingulum Mid Right |
| | | | Cingulum Post Right | Cingulum Post Left |
| | | | | Cingulum Post Right |
| **Temporal Lobe** | Heschl Right | Amygdala Left | Hippocampus Left | ParaHippocampal Left |
| | Temporal Sup Left | Heschl Left | ParaHippocampal Left | Amygdala Left |
| | Temporal Pole Sup Left | Heschl Right | ParaHippocampal Right | Temporal Sup Right |
| | Temporal Pole Mid Left | | Amygdala Left | Temporal Pole Sup Left |
| | | | Amygdala Right | Temporal Mid Left |
| | | | Temporal Pole Sup Left | Temporal Inf Right |
| | | | Temporal Pole Sup Right | Heschl Left |
| | | | Temporal Pole Mid Left | Heschl Right |
| | | | Temporal Pole Mid Right | |
| | | | Heschl Left | |

| | | | | |
|---|---|---|---|---|
| **Parietal Lobe** | Postcentral Left | Postcentral Left | Postcentral Right | Postcentral Left |
| | Postcentral Right | Postcentral Right | Parietal Sup Left | Postcentral Right |
| | Parietal Sup Left | Parietal Sup Left | Parietal Sup Right | Parietal Sup Right |
| | Parietal Sup Right | Parietal Sup Right | Parietal Inf Right | Parietal Inf Left |
| | Parietal Inf Left | Precuneus Right | Precuneus Right | Parietal Inf Right |
| | Parietal Inf Right | Paracentral Lobule Left | Paracentral Lobule Left | SupraMarginal Left |
| | SupraMarginal Left | Paracentral Lobule Right | Paracentral Lobule Right | SupraMarginal Right |
| | SupraMarginal Right | | | Precuneus Left |
| | Angular Right | | | Precuneus Right |
| | Precuneus Left | | | Paracentral Lobule Left |
| | Precuneus Right | | | Paracentral Lobule Right |
| | Paracentral Lobule Right | | | |
| **Occipital Lobe** | Occipital Sup Right | Calcarine Left | Calcarine Left | Calcarine Left |
| | Fusiform Right | Occipital Sup Left | Calcarine Right | Calcarine Right |
| | | Occipital Mid Left | Lingual Left | Lingual Left |
| | | Occipital Mid Right | Lingual Right | Lingual Right |
| | | Fusiform Left | Occipital Inf Right | Occipital Inf Left |
| | | | | Fusiform Left |
| | | | | Fusiform Right |
| **Subcortical structures** | Thalamus Left | Thalamus Left | Thalamus Left | Caudate Left |
| | Thalamus Right | Thalamus Right | Thalamus Right | Caudate Right |
| | | | | Putamen Right |
| | | | | Thalamus Left |
| | | | | Thalamus Right |

**Table S3.** AAL labels of the significant parcels ($p < .05$) found for the beta band during SMA stimulation for Go Frequent and Go Rare groups for action preparation trials (i.e., -700 ms SOA, -300 ms SOA).

| BETA BAND (13-30 Hz) | IFG | | | |
|---|---|---|---|---|
| | GO Frequent | | GO Rare | |
| | -700 | -300 | -700 | -300 |
| **Frontal Lobe** | Rolandic Oper Right | Rolandic Oper Right | Precentral Left | Precentral Left |
| | Cingulum Post Left | Cingulum Post Right | Frontal Sup Right | Precentral Right |
| | Cingulum Post Right | | Frontal Mid Left | Frontal Sup Left |
| | | | Frontal Mid Right | Frontal Sup Right |
| | | | Frontal Mid Orb Left | Frontal Mid Right |
| | | | Frontal Inf Tri Right | Frontal Mid Orb Right |
| | | | Frontal Inf Orb Right | Supp Motor Area Left |
| | | | Rolandic Oper Left | Supp Motor Area Right |
| | | | Frontal Med Orb Right | Olfactory Right |
| | | | Cingulum Post Right | Insula Right |
| | | | | Cingulum Ant Left |
| | | | | Cingulum Ant Right |
| **Temporal Lobe** | Heschl Right | Hippocampus Right | Amygdala Left | Hippocampus Right |
| | Temporal Sup Right | ParaHippocampal Right | Heschl Right | Amygdala Right |
| | Temporal Inf Left | | Temporal Sup Right | Heschl Left |
| | | | Temporal Pole Sup Left | Temporal Sup Right |
| | | | Temporal Pole Sup Right | |
| | | | Temporal Pole Mid Right | |
| | | | Temporal Inf Right | |
| **Parietal Lobe** | Postcentral Right | Parietal Sup Left | Parietal Sup Right | Postcentral Left |
| | Parietal Sup Right | Precuneus Left | SupraMarginal Right | Precuneus Right |
| | | Paracentral Lobule Right | Precuneus Right | |
| **Occipital Lobe** | Calcarine Right | Calcarine Left | Calcarine Left | Calcarine Left |
| | Cuneus Right | Cuneus Left | Calcarine Right | Calcarine Right |
| | Occipital Sup Left | Lingual Right | Cuneus Left | Cuneus Left |
| | Occipital Sup Right | Occipital Sup Left | Cuneus Right | Cuneus Right |
| | Occipital Inf Left | Occipital Sup Right | Occipital Sup Left | Occipital Inf Left |

| | | | | |
|---|---|---|---|---|
| | Occipital Inf Right | Occipital Inf Right | Occipital Inf Right | Occipital Inf Right |
| | | Fusiform Right | | Fusiform Left |
| **Subcortical Structures** | | | Thalamus Right | Putamen Right |

**Table S4.** AAL labels of the significant parcels ($p < .05$) found for the beta band during IFG stimulation for Go Frequent and Go Rare groups for action preparation trials (i.e., -700 ms SOA, -300 ms SOA).

| ALPHA BAND (8-12 Hz) | SMA | | IFG | |
|---|---|---|---|---|
| | **GO trial** | **No-Go trial** | **Go trial** | **No-Go trial** |
| **Frontal lobe** | Precentral Left | Frontal Sup Left | Precentral Left | Precentral Left |
| | Frontal Sup Orb Left | Frontal Mid Left | Frontal Sup Left | Frontal Sup Right |
| | Frontal Sup Orb Right | Frontal Mid Right | Frontal Mid Left | Frontal Sup Orb Left |
| | Frontal Mid Left | Frontal Inf Oper Right | Supp Motor Area Left | Frontal Sup Orb Right |
| | Frontal Mid Right | Rolandic Oper Right | Supp Motor Area Right | Frontal Mid Left |
| | Frontal Mid Orb Left | Supp Motor Area Right | Insula Left | Frontal Mid Right |
| | Frontal Inf Oper Right | Frontal Sup Medial Left | Insula Right | Frontal Mid Orb Left |
| | Frontal Inf Tri Left | Cingulum Ant Left | Cingulum Mid Left | Frontal Mid Orb Right |
| | Frontal Inf Tri Right | Cingulum Mid Left | Cingulum Mid Right | Frontal Inf Orb Left |
| | Rolandic Oper Left | Cingulum Mid Right | Cingulum Post Right | Frontal Inf Orb Right |
| | Supp Motor Area Right | | | Rolandic Oper Right |
| | Olfactory Left | | | Supp Motor Area Left |
| | Olfactory Right | | | Olfactory Left |
| | Frontal Med Orb Left | | | Olfactory Right |
| | Frontal Med Orb Right | | | Frontal Sup Medial Left |
| | Rectus Left | | | Frontal Sup Medial Right |
| | Rectus Right | | | Frontal Med Orb Right |
| | Insula Left | | | Rectus Right |
| | Cingulum Ant Left | | | Insula Left |
| | Cingulum Ant Right | | | Insula Right |
| | Cingulum Mid Left | | | Cingulum Ant Left |
| | Cingulum Mid Right | | | Cingulum Ant Right |
| | Cingulum Post Right | | | Cingulum Mid Left |
| | | | | Cingulum Mid Right |
| | | | | Cingulum Post Left |
| | | | | Cingulum Post Right |
| **Temporal Lobe** | Hippocampus Left | Hippocampus Right | Temporal Mid Right | Hippocampus Left |
| | Hippocampus Right | ParaHippocampal Right | Temporal Inf Right | Hippocampus Right |
| | ParaHippocampal Right | Heschl Left | | ParaHippocampal Left |

| | | | | |
|---|---|---|---|---|
| | Amygdala Left | Heschl Right | | Amygdala Left |
| | Amygdala Right | Temporal Sup Right | | Amygdala Right |
| | Heschl Left | Temporal Pole Sup Left | | Heschl Left |
| | Heschl Right | Temporal Pole Sup Right | | Heschl Right |
| | Temporal Sup Left | Temporal Pole Mid Left | | Temporal Sup Left |
| | Temporal Sup Right | Temporal Pole Mid Right | | Temporal Sup Right |
| | Temporal Mid Left | Temporal Inf Left | | Temporal Pole Sup Left |
| | Temporal Mid Right | Temporal Inf Right | | Temporal Pole Sup Right |
| | Temporal Pole Mid Right | Temporal Pole Mid Left | | Temporal Mid Left |
| | Temporal Inf Right | Temporal Pole Mid Right | | Temporal Mid Right |
| | | | | Temporal Pole Mid Left |
| | | | | Temporal Pole Mid Right |
| | | | | Temporal Inf Left |
| | Postcentral Left | Postcentral Left | Postcentral Left | Postcentral Left |
| | Postcentral Right | Parietal Sup Left | Postcentral Right | Parietal Inf Left |
| | Parietal Inf Left | Parietal Sup Right | Parietal Sup Left | SupraMarginal Right |
| | Parietal Inf Right | Parietal Inf Left | Parietal Sup Right | Angular Left |
| | Angular Right | Parietal Inf Right | Parietal Inf Left | Angular Right |
| **Parietal Lobe** | Precuneus Left | SupraMarginal Left | Parietal Inf Right | Precuneus Left |
| | Precuneus Right | SupraMarginal Right | SupraMarginal Right | Precuneus Right |
| | | Angular Left | Angular Right | Paracentral Leftobule Left |
| | | Angular Right | Precuneus Left | Paracentral Leftobule Right |
| | | | Precuneus Right | |
| | | | Paracentral Leftobule Left | |
| | Occipital Sup Right | Calcarine Left | Calcarine Left | Calcarine Left |
| | Occipital Mid Right | Cuneus Left | Calcarine Right | Calcarine Right |
| | Occipital Inf Right | Cuneus Right | Cuneus Right | Cuneus Left |
| **Occipital Lobe** | | Occipital Sup Left | Lingual Left | Cuneus Right |
| | | Occipital Sup Right | Occipital Sup Left | Lingual Left |
| | | Occipital Mid Right | Occipital Sup Right | Lingual Right |
| | | Occipital Inf Left | Occipital Mid Left | Occipital Sup Left |

| | | | | |
|---|---|---|---|---|
| | | Occipital Inf Right | Occipital Mid Right | Occipital Sup Right |
| | | Fusiform Left | Occipital Inf Left | Occipital Mid Left |
| | | | Occipital Inf Right | Occipital Mid Right |
| | | | | Occipital Inf Left |
| | | | | Occipital Inf Right |
| | | | | Fusiform Left |
| | | | | Fusiform Right |
| **Subcortical Structures** | Caudate Left | Thalamus Left | Putamen Right | Caudate Left |
| | Caudate Right | Thalamus Right | | Caudate Right |
| | Putamen Right | | | Putamen Right |
| | Thalamus Left | | | Thalamus Left |
| | Thalamus Right | | | Thalamus Right |

**Table S5.** AAL labels of the significant parcels ($p < .05$) found for the alpha band during SMA and IFG stimulation for Go trials and No-Go trials for +200 SOA.

| BETA BAND (13-30 Hz) | SMA | | IFG | |
|---|---|---|---|---|
| | **GO trial** | **No-Go trial** | **Go trial** | **No-Go trial** |
| **Frontal Lobe** | Precentral Left | Precentral Left | | Precentral Left |
| | Precentral Right | Precentral Right | | Precentral Right |
| | Frontal Sup Left | Frontal Sup Left | | Frontal Sup Orb Left |
| | Frontal Sup Orb Left | Frontal Sup Orb Left | | Frontal Sup Orb Right |
| | Frontal Mid Left | Frontal Sup Orb Right | | Frontal Mid Left |
| | Frontal Mid Right | Frontal Mid Left | | Frontal Mid Right |
| | Frontal Mid Orb Left | Frontal Mid Orb Left | | Frontal Mid Orb Left |
| | Frontal Inf Oper Left | Frontal Mid Orb Right | | Rolandic Oper Left |
| | Frontal Inf Tri Left | Frontal Inf Oper Right | | Olfactory Right |
| | Frontal Inf Tri Right | Frontal Inf Orb Left | | Frontal Sup Medial Left |
| | Frontal Inf Orb Left | Rolandic Oper Right | | Frontal Sup Medial Right |
| | Rolandic Oper Left | Supp Motor Area Right | | Rectus Right |
| | Insula Left | Olfactory Left | | Insula Left |
| | Cingulum Ant Right | Olfactory Right | | Insula Right |
| | Cingulum Mid Left | Frontal Sup Medial Left | | Cingulum Ant Left |
| | Cingulum Mid Right | Frontal Sup Medial Right | | Cingulum Post Left |
| | Cingulum Post Left | Frontal Med Orb Left | | |
| | Cingulum Post Right | Frontal Med Orb Right | | |
| | | Rectus Left | | |
| | | Rectus Right | | |
| | | Insula Left | | |
| | | Insula Right | | |
| | | Cingulum Ant Left | | |
| | | Cingulum Ant Right | | |
| | | Cingulum Mid Left | | |
| | | Cingulum Mid Right | | |
| | | Cingulum Post Left | | |
| | | Cingulum Post Right | | |
| **Temporal Lobe** | Hippocampus Right | Hippocampus Left | Hippocampus Right | Hippocampus Left |

| | | | | |
|---|---|---|---|---|
| | ParaHippocampal Right | Hippocampus Right | ParaHippocampal Right | Hippocampus Right |
| | Amygdala Left | ParaHippocampal Right | Temporal Inf Left | ParaHippocampal Left |
| | Amygdala Right | Amygdala Left | | Amygdala Left |
| | Heschl Left | Amygdala Right | | Amygdala Right |
| | Temporal Sup Left | Heschl Left | | Heschl Left |
| | Temporal Pole Sup Left | Temporal Pole Sup Left | | Temporal Sup Left |
| | Temporal Mid Right | Temporal Mid Right | | Temporal Pole Sup Right |
| | Temporal Pole Mid Left | Temporal Pole Mid Left | | Temporal Mid Left |
| | Temporal Pole Mid Right | Temporal Pole Mid Right | | Temporal Pole Mid Left |
| | | Temporal Inf Right | | Temporal Pole Mid Right |
| | | | | Temporal Inf Left |
| **Parietal Lobe** | Postcentral Left | Postcentral Left | Postcentral Right | Postcentral Left |
| | Parietal Sup Right | Postcentral Right | Parietal Sup Right | Postcentral Right |
| | Parietal Inf Right | Parietal Sup Left | Parietal Inf Right | Parietal Sup Left |
| | Angular Left | Parietal Sup Right | Paracentral Lobule Left | Parietal Inf Right |
| | Angular Right | Parietal Inf Left | Paracentral Lobule Right | Precuneus Right |
| | Precuneus Right | Parietal Inf Right | | Paracentral Lobule Left |
| | Paracentral Lobule Left | SupraMarginal Left | | Paracentral Lobule Right |
| | | SupraMarginal Right | | |
| | | Angular Left | | |
| | | Angular Right | | |
| | | Precuneus Left | | |
| | | Precuneus Right | | |
| | | Paracentral Lobule Left | | |
| | | Paracentral Lobule Right | | |
| **Occipital Lobe** | Cuneus Right | Calcarine Left | Calcarine Left | Calcarine Left |
| | Lingual Right | Calcarine Right | Calcarine Right | Calcarine Right |
| | Occipital Sup Right | Cuneus Left | Cuneus Right | Cuneus Left |
| | Occipital Mid Right | Cuneus Right | Lingual Left | Cuneus Right |
| | | Lingual Left | Lingual Right | Occipital Sup Right |
| | | Lingual Right | Occipital Mid Left | Occipital Mid Left |

| | | | | |
|---|---|---|---|---|
| | | Occipital Sup Left | Fusiform Left | Occipital Inf Left |
| | | Occipital Sup Right | | Occipital Inf Right |
| | | Occipital Mid Left | | Fusiform Left |
| | | Occipital Mid Right | | |
| | | Occipital Inf Left | | |
| | | Occipital Inf Right | | |
| | | Fusiform Left | | |
| **Subcortical Structures** | Caudate Left | Caudate Left | Thalamus Left | Putamen Right |
| | Caudate Right | Caudate Right | | Thalamus Right |
| | Thalamus Left | Putamen Right | | |
| | Thalamus Right | Thalamus Left | | |
| | | Thalamus Right | | |

**Table S6.** AAL labels of the significant parcels ($p < .05$) found for the beta band during SMA and IFG stimulation for Go trials and No-Go trials for +200 SOA.
